# MKFGO: Integrating Multi-Source Knowledge Fusion with Pre-Trained Language Model for High-Accuracy Protein Function Prediction

**DOI:** 10.1101/2025.03.27.645685

**Authors:** Yi-Heng Zhu, Shuxin Zhu, Xuan Yu, He Yan, Yan Liu, Xiaojun Xie, Dong-Jun Yu, Rui Ye

## Abstract

Accurately identifying protein functions is essential to understand life mechanisms and thus advance drug discovery. Although biochemical experiments are the gold standard for determining protein functions, they are often time-consuming and labor-intensive. Here, we proposed a novel composite deep-learning method, MKFGO, to infer Gene Ontology (GO) attributes through integrating five complementary pipelines built on multi-source biological data. MKFGO was rigorously benchmarked on 1522 non-redundant proteins, demonstrating superior performance over 11 state-of-the-art function prediction methods. Comprehensive data analyses revealed that the major advantage of MKFGO lies in its two deep-learning components, HFRGO and PLMGO, which derive handcraft features and protein large language model (PLM)-based features, respectively, from protein sequences in different biological views, with effective knowledge fusion at the decision-level. HFRGO leverages an LSTM-attention network embedded with handcraft features, in which the triplet loss-based guilt-by-association strategy is designed to enhance the correlation between feature similarity and function similarity. PLMGO employs the PLM to capture feature embeddings with discriminative functional patterns from sequences. Meanwhile, another three components provide complementary insights for further improving prediction accuracy, driven by protein-protein interaction, GO term probability, and protein-coding gene sequence, respectively. The source codes and models of MKFGO are freely available at https://github.com/yiheng-zhu/MKFGO.

## INTRODUCTION

Proteins play a fundamental role in various biological processes, such as enzymatic activity, gene expression regulation, and supporting cell structures [1, 2]. Accurate identification of protein functions is vital to unravel life mechanisms and guide drug design, with functions categorized into three aspects, i.e., molecular function (MF), biological process (BP), and cellular component (CC), under the widely used Gene Ontology (GO) annotation [3, 4]. While biochemical experiments are the gold standard for determining protein functions, they are often labor-intensive and may yield incomplete results, leaving numerous sequenced proteins without known functional annotations [5]. As of March 2025, the UniProt database [6] housed ∼253 million protein sequences, but fewer than 0.1% were annotated with GO terms supported by experimental evidence. To bridge this gap, there is an urgent need to develop efficient computational methods for protein function prediction [7, 8].

The existing function prediction methods can be divided into three categories: template detection, statistical machine learning, and deep learning-based methods. In the early stage, template detection-based methods were predominant in function prediction, focusing on identifying templates with similar sequences or structures to the query for functional inference [9, 10]. For example, GoFDR [11] and Blast2GO [12] utilize BLAST alignments [13] to search sequence templates, whereas FINDSITE [14] and COFACTOR [15] employ TM-align [16] to detect structure templates.

An inherent drawback of template detection-based methods is that their prediction accuracy heavily depends on the availability and quality of functional templates. To eliminate this dependence, statistical machine learning algorithms have been employed as an alternative. This could be implemented by extracting handcraft feature representations (e.g., k-mer sequence encoding [17] and position-specific scoring matrix [18]) from protein sequences, which are then processed with statistical machine learning algorithms (e.g., support vector machine [19] and random forest [20]) to train function prediction models, exemplified by GOPred [21], FFPred [22], and GOlabeler [23]. Although these methods complement template detection methods, their prediction accuracy is still insufficient [24]. The major reason is that the used machine learning models fail to derive the deep-level functional patterns buried in feature representations. To partly address these issues, deep learning techniques have emerged in function prediction [25].

The significant advantage of deep learning methods over statistical machine learning methods is that they can capture the sophisticated functional patterns from handcrafted feature representations through designing complex neural networks, such as convolution neural network (CNN) [26] and long short-term memory (LSTM) [27], with the classical examples of DeepGO [25], DeepGOCNN [28], TALE [29], DeepGOZero [30], and AnnoPRO [31]. Moreover, the protein large language models (PLMs), such as ESM2 [32] and ProtTrans [33], are increasingly demonstrating their potential in the feature representation through encoding the primary sequences as the discriminative feature embeddings. Since the PLMs are pre-trained on the networks with dozens of layers over hundreds of millions of sequences, they learn the abundant evolution knowledge related to function, resulting in the encoded feature embeddings containing distinctive functional patterns. Therefore, several function prediction methods directly employ PLMs to generate feature embeddings instead of traditional handcraft features, which are then fed to neural networks for implementing prediction models. The typical examples include ATGO [34], GAT-GO [35], SPROF-GO [36], TransFew [37], and DeepGO-SE [38].

Despite the great progress, challenges remain. First, the above-mentioned works have entirely replaced handcrafted features with PLM-based features, potentially leading to the incomplete capture of functional patterns. The underlying reason is that PLM-based features focus solely on the view of protein sequence evolution, whereas handcrafted features can extract function-related knowledge from other complementary views, such as protein secondary structure and family. Thus, the effective fusion of the knowledge from handcrafted and PLM-based features remains a significant challenge. Second, most existing function prediction methods derive knowledge from sequence alone, overlooking other crucial biological data sources (e.g., protein-protein interaction network and protein-coding gene) that contain complementary knowledge. Therefore, another challenge lies in the integration of multiple biological data sources to further improve prediction performance.

In this work, we proposed MKFGO, a composite protein function prediction method through integrating five complementary pipelines built on multi-source biological data. First, we designed two deep learning-based GO prediction pipelines, HFRGO and PLMGO, embedded with handcrafted and PLM-based features, respectively, from amino acid sequences. HFRGO leverages the LSTM-attention architecture with three powerful handcrafted features from the views of sequence conversion, secondary structure, and family domain, respectively. Meanwhile, the triplet loss-based guilt-by-association strategy [34] is employed to enhance the correlation between sequence feature similarity and function similarity. PLMGO employs the ProtTrans transformer [33] to encode the sequences as feature embeddings with functional patterns from the view of evolution diversity, which are then decoded by the fully connected neural network. Second, we implemented another three pipelines, driven by protein-protein interaction inference, naïve probability, and coding-gene sequence. Finally, a composite model was derived by incorporating the outputs of five complementary pipelines. Computational experiments on 1522 non-redundant test proteins have demonstrated two points. First, HFRGO and PLMGO complement each other, with decision-level knowledge fusion outperforming feature-level fusion. Second, the composite MKFGO exhibits a significant advantage in the accurate prediction of GO terms over the existing state-of-the-art approaches, as each of its five components contributes to the overall performance improvement. The source codes and models of MKFGO are freely available at https://github.com/yiheng-zhu/MKFGO.

## MATERIALS AND METHODS

### Benchmark datasets

We employed an approach closely mirroring the Critical Assessment of protein Function Annotation (CAFA) experiment to construct benchmark datasets. Specifically, we downloaded all protein sequences from the UniProt database [6] with the corresponding functional annotations from the Gene Ontology Annotation (GOA) database [39]. Then, we filtered out proteins through only selecting those that have been manually reviewed with the available function annotations by at least one of the eight experimental evidence codes, including EXP, IDA, IPI, IMP, IGI, IEP, TAS, and IC [40, 41]. After this, we collected 80653 high-quality proteins, which could be further split into training, validation, and test datasets. The 1522 proteins were selected as the test datasets, which were released in the UniProt database after 2021-07-01; and the 974 proteins as the validation datasets, released in the UniProt from 2020-07-01 to 2021-06-30. The remaining proteins have been filtered out by removing the redundant proteins aligned with test and validation proteins using CD-HIT [42] software with a sequence identity cut-off of 30%, yielding a training dataset of 70712 proteins.

The numbers of entries in each dataset across different GO categories are presented in **Table S1** of Supporting Information (SI). The training, validation, and test datasets were used independently to train models, optimize the models’ parameters, and assess the models’ performance.

### The architecture of MKFGO

As depicted in **Figure 1**, MKFGO is a composite deep-learning model for protein function prediction, where the input is a protein sequence with UniProt ID, and the output includes the confidence scores of predicted functional terms for three GO aspects. This model consists of five pipelines, i.e., (**A**) handcraft feature representation-based GO prediction (HFRGO), (**B**) protein language model-based GO prediction (PLMGO), (**C**) protein-protein interaction-based GO prediction (PPIGO), (**D**) naïve-based GO prediction (NAIGO), and (**E**) DNA language model-based GO prediction (DLMGO), which are driven by the protein sequence (**A** and **B**), interaction network (**C**), GO term probability (**D**), and coding-gene sequence (**E**), respectively. The input sequence is independently fed to five pipelines to generate the confidence scores of GO terms, which are further ensembled by the neural network to output the consensus scores.

**Figure 1.**
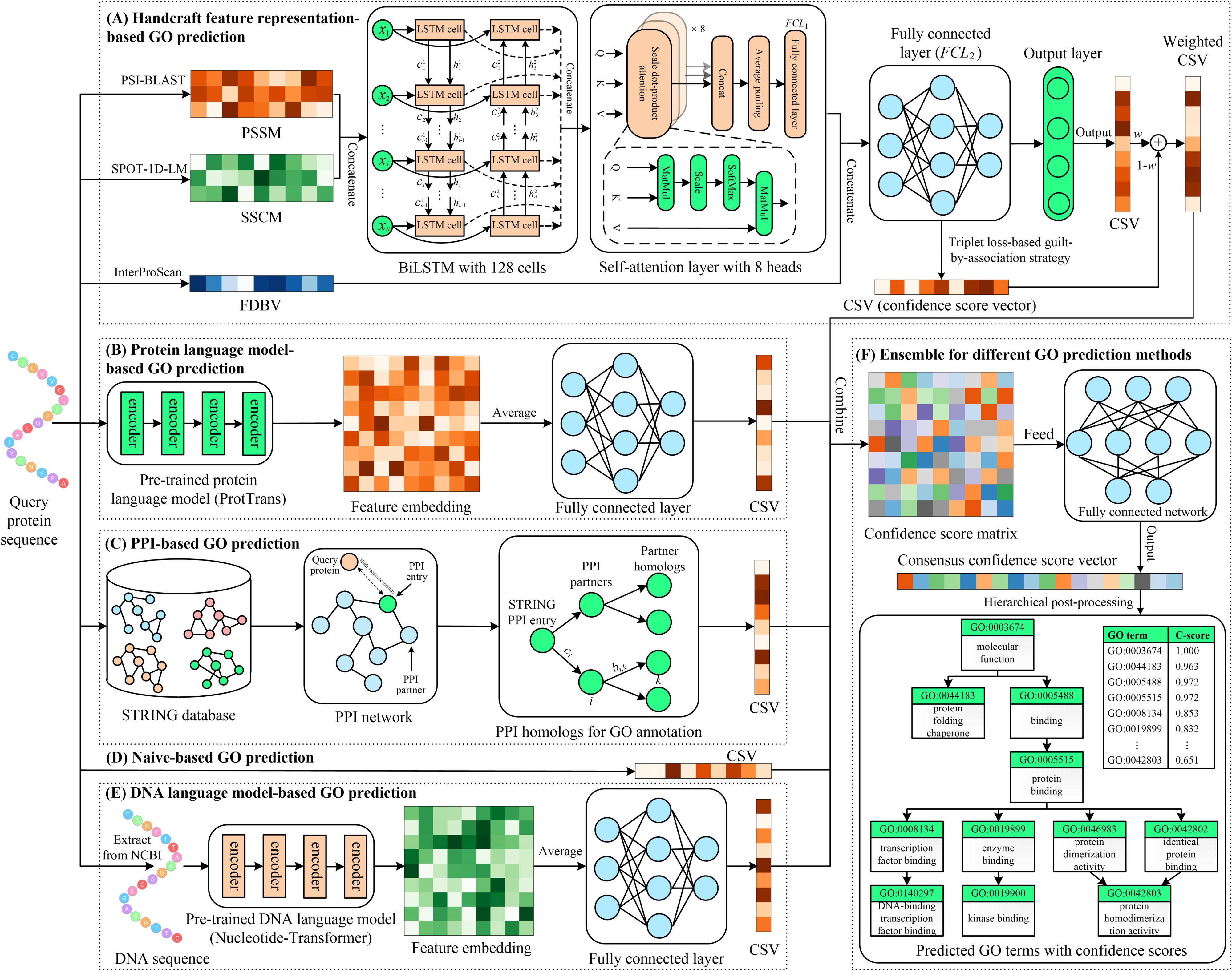
The flowchart of MKFGO.

### (A) Handcraft feature representation-based GO prediction

#### (I) Feature representation

For the query sequence with the length *L*, we use PSI-BLAST [13], SPOT-1D-LM [43], and InterProScan [44] programs to extract the corresponding feature representations, i.e., position-specific scoring matrix (PSSM), secondary structure coding matrix (SSCM), and family domain-based binary vector (FDBV), with the scale of *L* × 20, *L*× 8, and 45899-D, respectively, see details in **Text S1** of SI.

#### (II) LSTM-attention network-based function prediction

The PSSM and SSCM are concatenated and then fed to an LSTM-attention module consisting of a BiLSTM layer with 128 cells, a self-attention layer with 8 heads, an average-pooling layer, and a fully connected layer (*FCL*_1_) with 1024 neurons. The output of this module is concatenated with the FDBV and then processed by another fully connected layer (*FCL*_2_) with 1024 neurons to output a feature embedding vector, as carefully described in **Text S2** of SI.

The feature embedding vector is further fed to the output layer with a Sigmoid activation function [45] to generate a confidence score vector ***s****_sig_* for predicted GO terms. Meanwhile, the triplet loss-based guilt-by-association (TL-GBA) strategy [34] is performed on this embedding vector to produce another confidence score vector ***s****_gba_*. Finally, two confidence score vectors are weightedly combined to generate the final confidence score vector ***s****_hfr_* for the HFRGO pipeline.

#### (III) Triplet loss-based guilt-by-association strategy

For a query protein, we select the top *K* templates, which have the highest sequence feature similarity with itself, from the training dataset for function annotation:

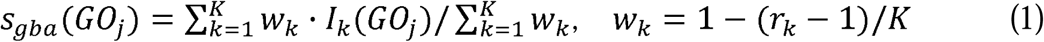

where *GO_j_* is the *j*-th candidate GO term; *I_k_*(*GO_j_*) = 1, if the *k*-th template is associated with *GO_j_* in the experimental annotation; otherwise, *I_k_*(*GO_j_*) = 0; *r_k_* is the rank of the *k*-th template in *K* templates based on the feature similarity with query.

The feature similarity between the template and query is measured by the Euclidean distance [46] of feature embedding vectors outputted by the *FCL*_2_. To improve the quality of selected templates, we employ the triplet loss [47] to enhance the correlation between sequence feature similarity and functional similarity:

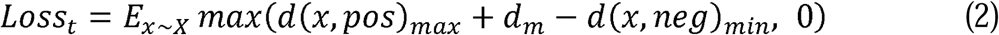

where *x* is a protein sequence in the training dataset *X*; *d_m_* is a pre-set margin value; *d*(*x, pos*)*_max_* (*d*(*x, neg*)*_min_*) is the maximum (minimum) values of distances between *x* and all positive (negative) partners which have the same (different) function to *x*. Two proteins are defined as having the same function if their functional similarity is higher than a pre-set threshold *c_f_*, as carefully described in **Text S3**. Minimizing this triplet loss helps ensure that the selected templates exhibit both higher feature similarity and functional similarity to the query.

#### (IV) Loss function

Considering that triplet loss is hardly converged in the training stage, we have added the cross-entropy loss to form a composite loss function [48-50]:

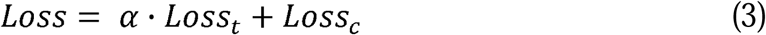

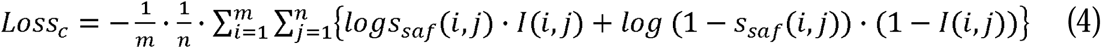

where *α* is a balanced parameter, *m* and *n* are the numbers of training proteins and GO terms, respectively; *s_saf_*(*i, j*) is the confidence score that the *i*-th training protein is associated with the *j*-th GO term predicted by the output layer in LSTM-attention network; *I*(*i, j*) = 1 if i-th protein is associated with the j-th term in the experimental annotation. This loss function could be minimized to optimize the hyper-parameters of the HFRGO using the Adam optimization algorithm [51]. In addition, the values of *d_m_, c_f_*, *α*, and *K* for GO aspects are listed in **Table S2** of SI.

### (B) Protein language model-based GO prediction

The input sequence is fed to the pre-trained protein language model, i.e., ProtTrans [33], to extract the feature embedding matrix, which is then averaged in the full sequence length to generate the embedding vector with 1024 dimensions. This vector is further processed by a fully connected layer with 1024 neurons and an output layer to yield the confidence score vector for GO terms. The cross-entropy loss (see details in **Equation 4**) is employed to optimize the hyper-parameters of the neural network. Here, the version of the ProtTrans model is ProtT5-XL-UniRef50, composed of 24 attention layers with 3 billion hyper-parameters and trained on over 45 million protein sequences.

### (C) Protein-protein interaction-based GO prediction

The Blastp [13] program is utilized to hit a PPI entry *P_e_*, which has the highest sequence identity to the query protein, against the STRING database [52]. For each PPI partner of *P_e_*, the Blastp is employed again with the e-value of 0.1 to search the corresponding homologs from the training sequence dataset. These homology proteins are used as templates for GO annotations, as carefully described in **Text S4**.

### (D) Naïve-based GO prediction

The confidence score that the query is associated with a GO term could be directly assigned by the frequency of this term in the training dataset:

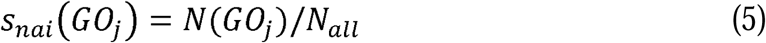

where *N*(*GO_j_*) and *N_all_* are the number of proteins associated with *GO_j_* and all proteins in the training dataset, respectively.

### (E) DNA language model-based GO prediction

For the query protein, we could download the DNA sequence of its coding gene from the NCBI [53] through mapping its UniProt ID to the Entrez ID of the coding gene. This DNA sequence is fed to the pre-trained DNA language model, i.e., Nucleotide-Transformer [54], to capture the feature embeddings, then further processed by the fully connected neural network to output the confidence score vector, using the same architecture in the PLMGO pipeline. Here, we employed two model versions of Nucleotide-Transformer, including NT-Multispecies (2.5B) and NT-1000G (2.5B), which are trained on more than 200 billion nucleotides from different species. Each language model could encode the DNA sequence as an embedding vector with 2560-D, which is then concatenated and fed to the neural network.

### (F) Ensemble for different GO prediction methods

The confidence score vectors of five GO prediction pipelines are concatenated as a confidence score matrix, which is then fed to a fully connected network layer with 512 neurons followed by an output layer with 1 neuron to output the consensus confidence scores of GO terms. Finally, a hierarchical post-processing procedure is performed on these confidence scores to ensure that the confidence score of a GO term is larger than or equal to those of its all children, as carefully described in **Text S5**.

### Evaluation metrics

Following the rules of CAFA competitions, we use three metrics to evaluate our models, i.e., maximum F_1_-score (F_max_), minimum semantic distance (S_min_), and area under the precision-recall curve (AUPRC) [55, 56]. F_max_ is the highest F-score achieved across all confidence thresholds, offering a single measure of the best trade-off between precision and recall. S_min_ measures the discrepancy between predicted and true GO terms by calculating the semantic distance in the GO hierarchy structure. AUPRC assesses a model’s overall performance in the trade-off between precision and recall over all thresholds. The detailed calculations of these metrics can be found in **Text S6**.

## RESULTS AND DISCUSSIONS

### Overall performance of MKFGO

We benchmarked the proposed methods with 11 state-of-the-art function prediction methods on all of 1522 test proteins, including eight single methods (Blast-KNN [23], FunFams [57], DeepGOCNN [28], TALE [29], DeepGOZero [30], ATGO [34], AnnoPRO [31], and DeepGO-SE [38]) and three composite methods (DeepGOPlus [28], TALE+ [29], and ATGO+ [34]). These single methods could be categorized into three groups: (1) Blast-KNN and FunFams are template detection-based methods, leveraging sequence homology alignment and protein family search separately; (2) DeepGOCNN, TALE, DeepGOZero, and AnnoPRO are deep learning-based methods with handcraft feature representations; (3) ATGO and DeepGO-SE are deep learning methods with PLM-based feature representations. Moreover, DeepGOPlus, TALE+, and ATGO+ are the composite versions for DeepGOCNN, TALE, and ATGO, respectively, through integrating Blast-KNN. Accordingly, our competing methods include the composite MKFGO and its three component methods (i.e., PPIGO, HFRGO, and PLMGO), each corresponding to one of the above-mentioned three groups.

**Table 1** summarizes the performance comparison between our methods and 11 existing methods on 1522 test proteins. Overall, the proposed MKFGO achieves the best performance among the 15 function prediction methods. In comparison to the second-best performer, i.e., ATGO+, MKFGO gains 4.8%, 5.3% [= (|6.97-7.22|/7.22 + |23.08-23.88|/23.88 + |7.38-8.11|/8.11)/3×100%], and 7.6% average improvement for F_max_, S_min_, and AUPRC, respectively, on three GO aspects. Moreover, the composite methods (i.e., ATGO+, TALE+, and DeepGOPlus) all exhibit superior performance compared to their individual deep-learning counterparts, as BLAST-KNN provides complementary knowledge for function prediction.

**Table 1.** The overall performance of 15 function prediction methods on all 1522 test proteins.

| Method | F<sub>max</sub> |  |  | S<sub>min</sub> |  |  | AUPRC |  |  | Coverage<sup>f</sup> |  |  |
| --- | --- | --- | --- | --- | --- | --- | --- | --- | --- | --- | --- | --- |
|  | MF | BP | CC | MF | BP | CC | MF | BP | CC | MF | BP | CC |
| Blast-KNN<sup>a, d</sup> | 0.642 | 0.397 | 0.485 | 7.77 | 24.90 | 8.59 | 0.346 | 0.220 | 0.259 | 0.832 | 0.803 | 0.717 |
| FunFams<sup>a, d</sup> | 0.483 | 0.311 | 0.387 | 9.87 | 27.24 | 9.02 | 0.298 | 0.141 | 0.200 | 0.631 | 0.599 | 0.532 |
| PPIGO<sup>a, d</sup> | 0.329 | 0.273 | 0.461 | 11.81 | 26.74 | 8.43 | 0.141 | 0.126 | 0.253 | 0.515 | 0.558 | 0.645 |
| DeepGOCNN<sup>b, e</sup> | 0.430 | 0.296 | 0.497 | 11.01 | 26.67 | 9.45 | 0.369 | 0.204 | 0.493 | 1.000 | 1.000 | 1.000 |
| TALE<sup>b, d</sup> | 0.457 | 0.313 | 0.526 | 11.19 | 25.88 | 8.77 | 0.397 | 0.222 | 0.534 | 1.000 | 1.000 | 1.000 |
| DeepGOZero<sup>b, d</sup> | 0.677 | 0.396 | 0.540 | 7.53 | 24.86 | 9.46 | 0.674 | 0.319 | 0.521 | 1.000 | 1.000 | 1.000 |
| AnnoPRO<sup>b, e</sup> | 0.504 | 0.365 | 0.535 | 9.63 | 25.36 | 8.67 | 0.366 | 0.267 | 0.504 | 1.000 | 1.000 | 1.000 |
| HFRGO<sup>b</sup> | 0.682 | 0.412 | 0.580 | 7.23 | 23.91 | 8.14 | 0.630 | 0.340 | 0.539 | 1.000 | 1.000 | 1.000 |
| ATGO<sup>c, d</sup> | 0.686 | 0.424 | 0.607 | 7.34 | 23.99 | 7.87 | 0.676 | 0.361 | 0.625 | 1.000 | 1.000 | 1.000 |
| DeepGO-SE<sup>c, d</sup> | 0.669 | 0.411 | 0.573 | 7.67 | 24.48 | 9.44 | 0.662 | 0.351 | 0.600 | 1.000 | 1.000 | 1.000 |
| PLMGO<sup>c</sup> | 0.680 | 0.424 | 0.628 | 7.58 | 23.95 | 7.57 | 0.621 | 0.355 | 0.571 | 1.000 | 1.000 | 1.000 |
| DeepGOPlus<sup>d</sup> | 0.660 | 0.402 | 0.574 | 7.78 | 24.92 | 8.56 | 0.620 | 0.311 | 0.517 | 1.000 | 1.000 | 1.000 |
| TALE+<sup>d</sup> | 0.640 | 0.401 | 0.581 | 8.04 | 24.91 | 8.37 | 0.617 | 0.318 | 0.550 | 1.000 | 1.000 | 1.000 |
| ATGO+<sup>d</sup> | 0.693 | 0.430 | 0.607 | 7.22 | 23.88 | 8.11 | 0.670 | 0.371 | 0.617 | 1.000 | 1.000 | 1.000 |
| MKFGO | **0.710** | **0.459** | **0.639** | **6.97** | **23.08** | **7.38** | **0.716** | **0.400** | **0.668** | 1.000 | 1.000 | 1.000 |

<sup>a</sup> Template detection-based methods; <sup>b</sup> Deep learning-based methods with handcraft feature representations; <sup>c</sup> Deep learning-based methods with PLM-based feature representations; <sup>d</sup> The prediction models are re-trained on our training dataset using the author's source codes; <sup>e</sup> The prediction models are directly downloaded from author's web platforms. <sup>f</sup> Coverage is the proportion of the number of test proteins with available prediction scores divided by the total number of test proteins. Bold fonts highlight the best performer in each category.

Among all single methods, our PLMGO and HFRGO are ranked 3/1/1 and 2/3/3 for MF/BP/CC aspects, respectively. Moreover, HFRGO outperforms all other deep learning methods that use handcrafted feature representations. Taking DeepGOZero as an example, our HFRGO beat it in 8 out of 9 evaluation metrics, except for the AUPRC in the MF aspect. Importantly, HFRGO consistently outperforms the DeepGO-SE, a PLM-based deep learning method, in terms of F_max_ and S_min_ values across three GO aspects. It is undeniable that ATGO achieves the highest prediction accuracy in MF aspects, likely because it utilizes PLM (i.e., the ESM-1b transformer [58]) to extract feature embeddings from three-level perspectives of sequence evolution, enriching the knowledge related to molecular functions.

Furthermore, we observe that deep learning methods, particularly those employing PLM, achieve significantly superior performance than template detection-based methods. Part of the reason is that such template detection methods cannot output any prediction results for some test proteins that fail to match available templates, leading to inferior performance in the overall test dataset with low coverage, especially evident in FunFams and PPIGO. Therefore, we conducted an additional benchmark of 15 function prediction methods on a subset of 515 test proteins, for which predictions can be produced by all methods. As illustrated in **Table S3**, a similar trend is observed, where our methods outperform the control methods by a substantial margin. Meanwhile, BLAST-KNN exhibits noticeably higher prediction accuracy over FunFams and PPIGO in both tests, indicating that sequence homology provides a more reliable basis for protein function inference than PPI and family similarity.

### MKFGO shows great generality to new species and non-homology proteins

Despite the progress in function prediction, many deep learning methods may exhibit reduced performance on proteins from new species absent from their training data. To assess the generalizability of MKFGO to new species, we mapped each protein in our dataset to its corresponding species and then gathered 300 test proteins from 158 new species that were never observed in the training dataset.

We further benchmarked the proposed MKFGO with 11 competing function prediction methods on these 300 new species proteins, where the corresponding F_max_ and S_min_ values across all GO aspects are shown in **Figure 2 (A-B)**. Meanwhile, the AUROC values of 12 methods are listed in **Figure S1**. It could be found that MKFGO achieves the best F_max_ and S_min_ values among all methods. Compared to the second-best performer, i.e., ATGO+, MKFGO gains an average improvement of 4.5% in F_max_ and 7.7% in S_min_, respectively, on three GO aspects. As for AUROC, MKFGO is ranked 2/3/1 for the MF/BP/CC aspect. Moreover, the AUROC gap between MKFGO and the top performer is minimal and almost negligible on the MF/BP aspect. Additionally, the F_max_, S_min_, and AUROC values of MKFGO for these 300 proteins in **Figure 2 (A-B)** are largely consistent with those of the entire test dataset in **Table 1**. These observations demonstrate that MKFGO maintains its strong performance when modeling new species proteins, highlighting the generalizability of its deep-learning approach.

**Figure 2.**
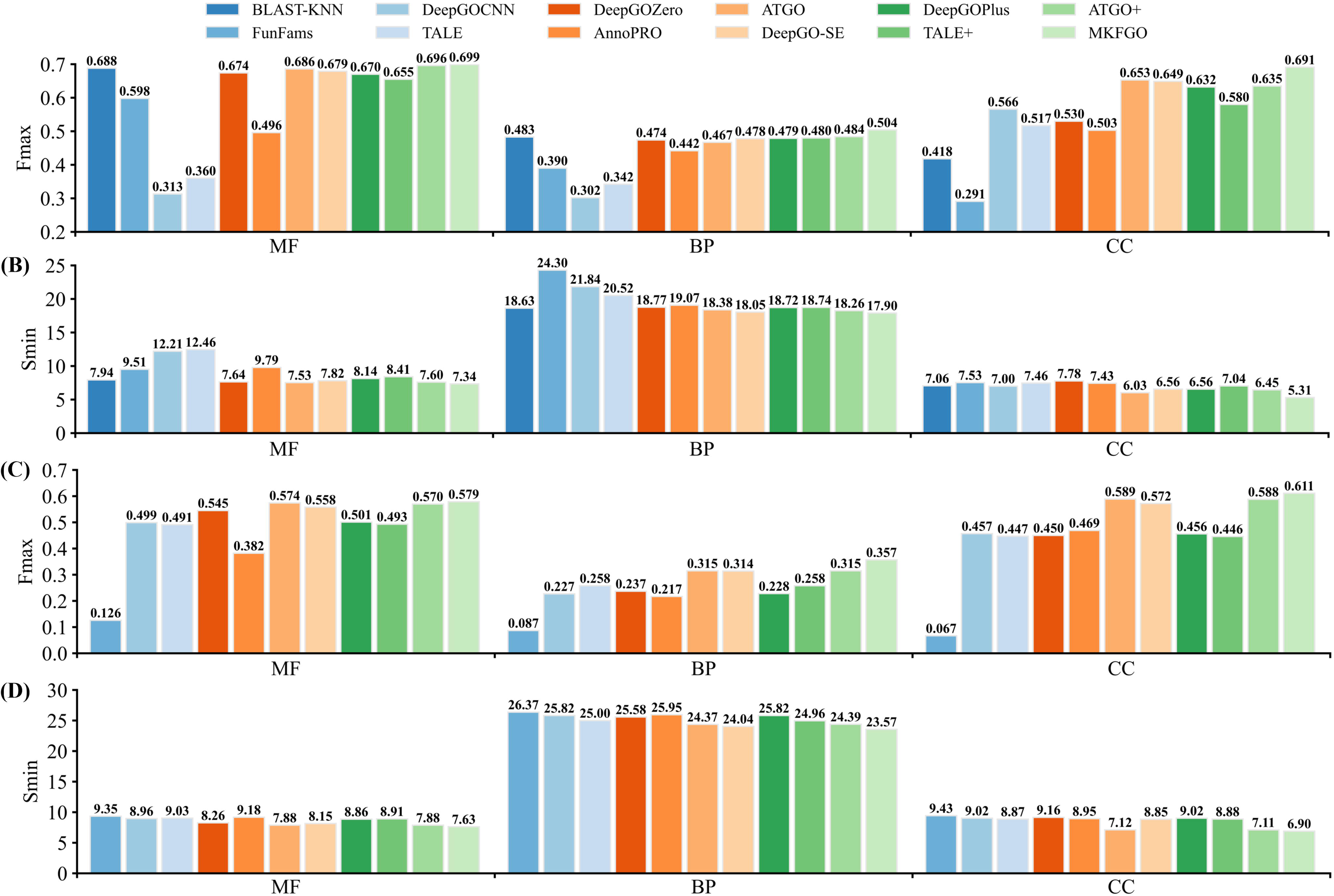
The performance comparison among 12 function prediction methods on the new species and non-homology proteins across three GO aspects. (A) The F_max_ values on the 300 test proteins from 158 new species. (B) The S_min_ values on the 300 new species proteins. (C) The F_max_ values on the 305 non-homology test proteins. (D) The S_min_ values on the 305 non-homology proteins.

Since the sequence-based function annotation is heavily dependent on sequence homology, another challenge for deep learning-based methods is the modeling of the proteins without sequence homologies. In light of this, we further benchmarked MKFGO with 10 competing methods on 305 test proteins that cannot hit any sequence homologies in the training dataset using BLAST search with an e-value of 0.01. Here, BLAST-KNN was excluded, because it cannot generate any predictions for these test proteins. **Figure 2 (C-D)** summarizes the F_max_ and S_min_ values of 11 function prediction methods for three GO aspects on 305 non-homology test proteins, where the corresponding AUROC values are illustrated in **Figure S2**. Overall, the performance of all prediction methods for these non-homology proteins in **Figure 2 (C-D)** is significantly inferior to that for the whole test dataset in **Table 1**. This observation further demonstrates the importance of sequence homology in function prediction both for template detection and deep learning-based methods. However, our MKFGO still outperforms the other 11 methods for all evaluation metrics across three GO aspects, except for the AUPRC value on the MF aspect. Taking ATGO+ as a reference, MKFGO achieves an improvement of 1.6%, 13.3%, and 3.9% on F_max_, and 3.2%, 3.4%, and 3.0% on S_min_ for MF, BP, and CC aspects, respectively.

### Ablation study

#### (A) Contribution analysis for different GO prediction components

To analyze the contributions of five component methods (i.e., HFRGO, PLMGO, PPIGO, NAIGO, and DLMGO) in MKFGO, we individually remove each component from the MKFGO to generate five reduced-composite methods, including PIND (PLMGO+PPIGO+NAIGO+DLMGO), HIND (HFRGO+PPIGO+NAIGO+DLMGO), HPND (HFRGO+PLMGO+NAIGO+DLMGO), HPID (HFRGO+PLMGO+PPIGO+ DLMGO), and HPIN (HFRGO+PLMGO+PPIGO+NAIGO). Here, “+” means that the component methods are ensembled using a fully connected neural network. We further benchmarked MKFGO with its component and reduced-composite methods on all 1522 test proteins, as summarized in **Figure 3**.

**Figure 3.**
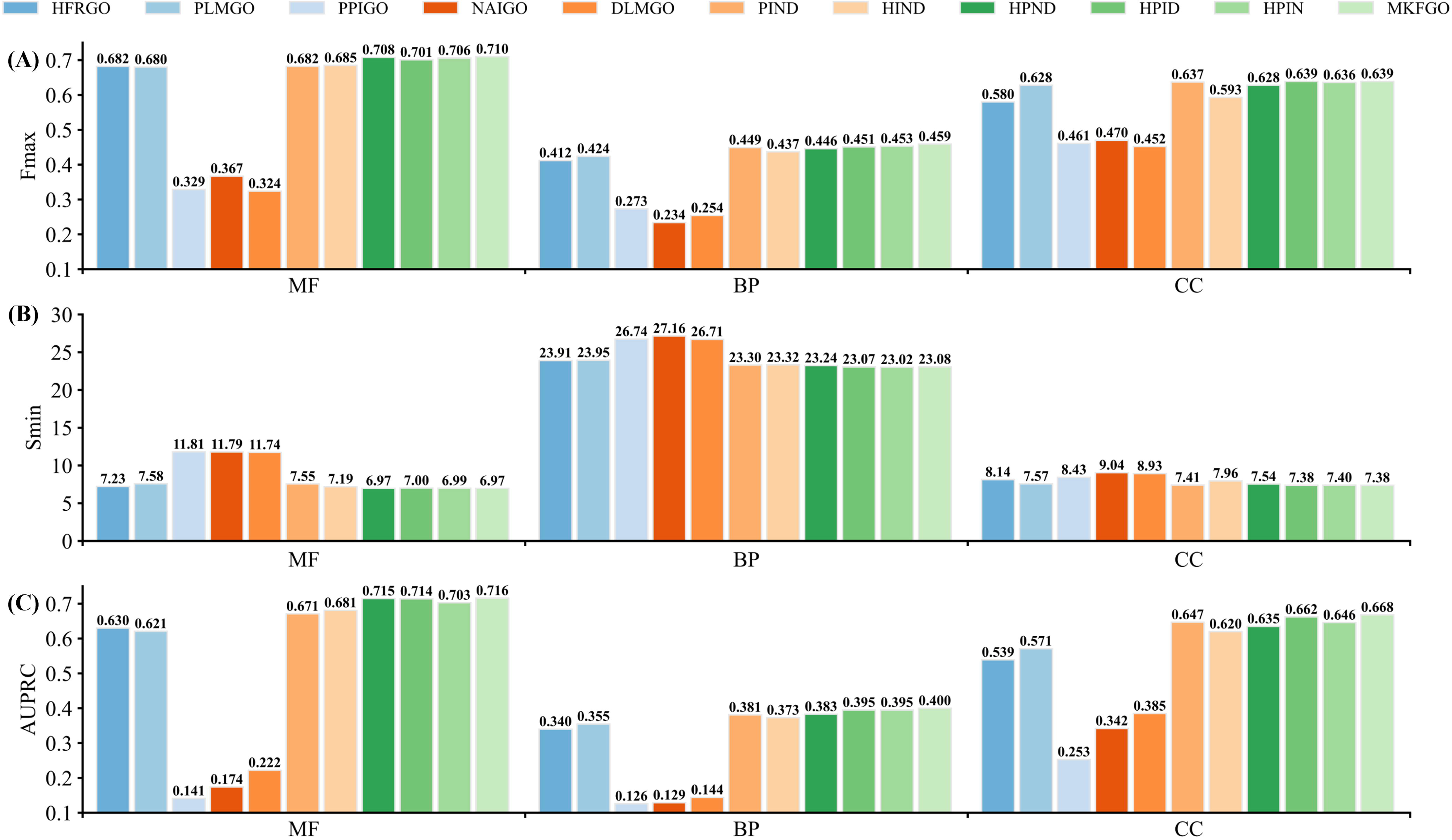
The performance comparison between MKFGO and its 10 component and reduced-composite methods on all 1522 test proteins across three GO aspects. (A) The F_max_ values. (B) The S_min_ values. (C) The AUPRC values.

It could be found that MKFGO yields the best performance across all evaluation metrics among the 11 prediction methods in the three GO aspects, except for the S_min_ value in the BP aspect. In terms of F_max_, for example, MKFGO shares an average increase of 2.2%, 5.5%, 1.6%, 1.0%, and 0.8%, respectively, on three GO aspects, in comparison to five reduced-composite methods, i.e., PING, HING, HPNG, HPIG, HPIN. This observation demonstrates that the five components both help to improve the prediction accuracy of MKFGO, indicating they could provide complementary knowledge for protein function prediction.

Among the five individual components, HFRGO and PLMGO are the top two performers, ranking 1/2/2 and 2/1/1 on MF/BP/CC aspects, respectively, with a small margin, on the balance of Fmax, Smin, and AUPRC values. Moreover, these two methods achieve a significant performance advantage over the other three components. After individually removing five components from MKFGO, the first and second largest performance decreases occur in HING and PING, with average decreases of 5.3% and 3.4%, respectively, for three evaluation metrics across all GO aspects. These data show that HFRGO and PLMGO make the most contributions to MKFGO, further demonstrating that the handcraft and PLM-based features from sequences have nearly equal efficacy for function prediction.

In MKFGO’s pipeline, we fused the knowledge buried in handcraft and PLM-based features at the decision-level rather than feature-level. The major reason is that incorporating too many features into a single neural network may lead to a learning bias toward certain features while overlooking other crucial ones. Moreover, such a network may not effectively handle the redundancy among multiple features. To demonstrate this point, we benchmarked the MKFGO with three designed control methods on 1522 test proteins, as shown in **Figure S3** and **Table S4** and discussed in **Text S7**. Experimental results further demonstrated that decision-level knowledge outperforms feature-level.

Additionally, the DLMGO method has the least contribution to MKFGO. This may be because of the limited correlation between the gene sequence and the function of the protein it encodes. However, another significant application of this method is predicting gene function from its DNA sequences, particularly for non-coding genes. Therefore, we constructed a benchmark dataset of genes, including a training set of 46084 protein-coding genes, a validation set of 144 non-coding genes, and a test set of 147 non-coding genes, from COXPRESdb [59] and ATTED-II [60] databases following the rules of TripletGO method [61], with the details in **Text S8** of SI. TripletGO is a composite gene function prediction method consisting of three components driven by gene expression data, gene sequence alignment, and naïve probability of GO terms, respectively. **Table S5** illustrates the performance comparison between DLMGO and TripletGO on 147 non-coning genes. The individual DLMGO demonstrates a performance comparable to the composite TripletGO. Moreover, integrating DLMGO as a component into TripletGO leads to an average improvement of 5.5%, 2.3%, and 13.5%, respectively, for F_max_, S_min_, and AUROC across three GO aspects. These results highlight the strong efficacy of DLMGO in non-coding gene function prediction.

#### (B) Performance comparison between different ensemble techniques

To examine the efficacy of the utilized fully connected neural network (FCNN) for integrating five components of MKFGO, we benchmarked it with three commonly used ensemble techniques, including logistic regression (LR), weighted voting (WV), and weighted product (WP), with the details in **Text S9** of SI. Specifically, for each GO term, the corresponding confidence scores of all components of MKFGO could be incorporated as a consensus score using one of the above four ensemble techniques.

Figure 4 illustrates the performance of four ensemble techniques on all 1522 test proteins. Our FCNN achieves the best performance among the four techniques, with an average 1.2% increase in F_max_ values compared to the second-best performer LR. Concerning S_min_ and AUPRC values, FCNN demonstrates superior performance to LR at least on two GO aspects. Regarding WV and WP, FCNN consistently outperforms across all evaluation metrics in all three GO aspects. It is worth noting that WP exhibited the poorest performance, even falling below that of the individual component method in **Table 1** on the MF aspect. This finding highlights the important role of the ensemble technique in composite function prediction.

**Figure 4.**
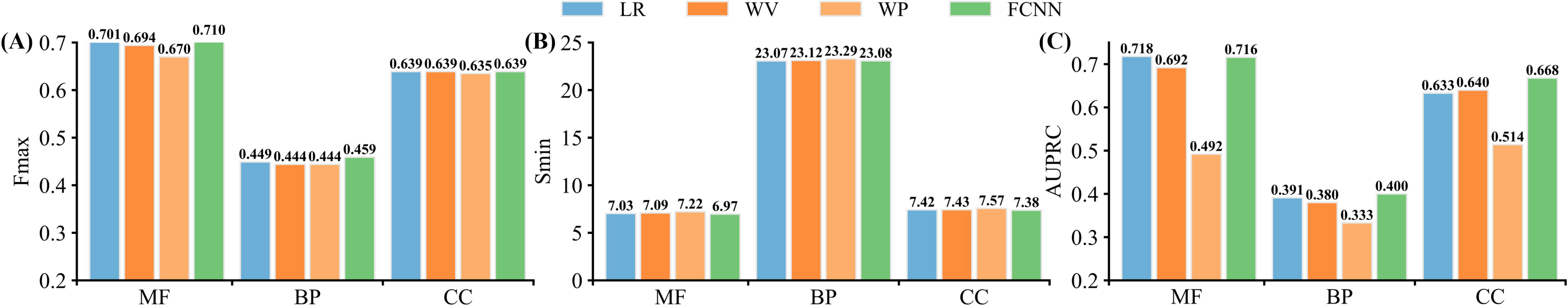
The performance comparison between four ensemble techniques for incorporating all components of MKFGO on all 1522 test proteins. (A) The F_max_ values. (B) The S_min_ values. (C) The AUPRC values.

#### (C) Ablation experiment for HFRGO

We conducted an ablation experiment to analyze the contributions of algorithmic innovations in HFRGO on its enhanced performance. Beginning with the HFRGO model (M0), we gradually remove algorithmic components. First, we remove the TL-GBA module (Module I) from M0 to build the model M1; Then, we individually exclude the [PSSM + SSCM + LSTM-attention layer] (Module II) and [FDBV + fully connected layer] (Module III) from M1 to develop the other two models (M2 and M3), with the architectures in **Figure S4**.

Figure 5 illustrates the performance comparison between four ablation models on all 1522 test proteins. Compared with M0, M1 without Module I exhibits reduced performance, with the average decrease of 1.0%, 1.2%, and 4.8% for F_max_, S_min_, and AUPRC, respectively, on three GO aspects. After individually removing Modules II and III from M1, the performance of M2 and M3 continuously drops. Taking M3 as an example, it shows inferior performance across all evaluation metrics except for the AUPRC values on MF and CC aspects, in comparison with M1. These data indicate that each of the three modules helps enhance the overall performance of HFRGO.

**Figure 5.**
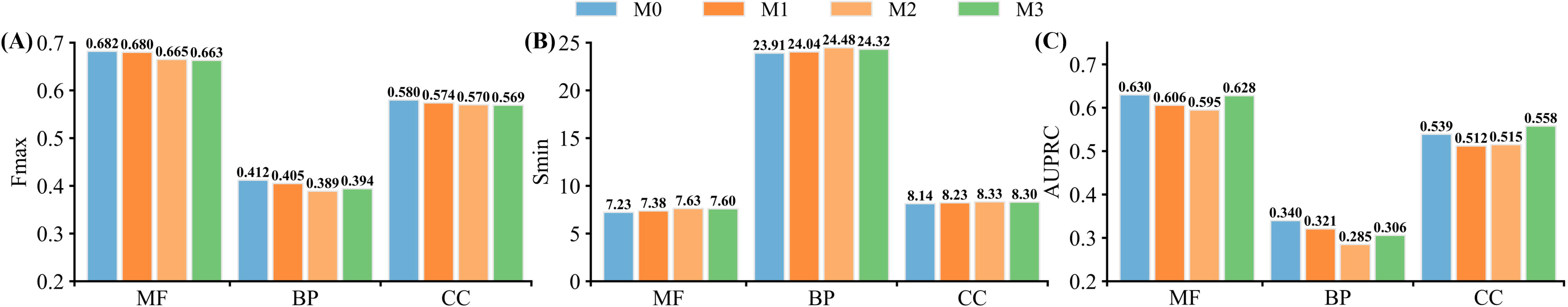
The performance comparison between four ablation models on all 1522 test proteins. (A) The F_max_ values. (B) The S_min_ values. (C) The AUPRC values.

### Case study

To further investigate the effects of different GO prediction methods, three representative proteins from our test dataset were selected for illustration, with the UniProt IDs of A0A2L2DDE6, Q8I2J3, and Q7Q2T8. These proteins are associated with 25, 14, and 7 GO terms, respectively, in the experimental annotation for the BP aspect, excluding the root term (GO:0008150, biological process). **Table 2** shows the performance comparison between MKFGO, its five components, and ATGO+ (i.e., the second-best performer in **Table 1**) on three representative proteins. Meanwhile, the correctly predicted GO terms (i.e., true positives) for these 7 methods are visualized as directed acyclic graphs in Figure 6. Moreover, the mistakenly predicted terms (false positives) for each method are listed in **Table S6**. It is worth noting that the predicted GO terms for different methods are determined by their respective cut-off setting to maximize the F_1_-score.

**Figure 6.**
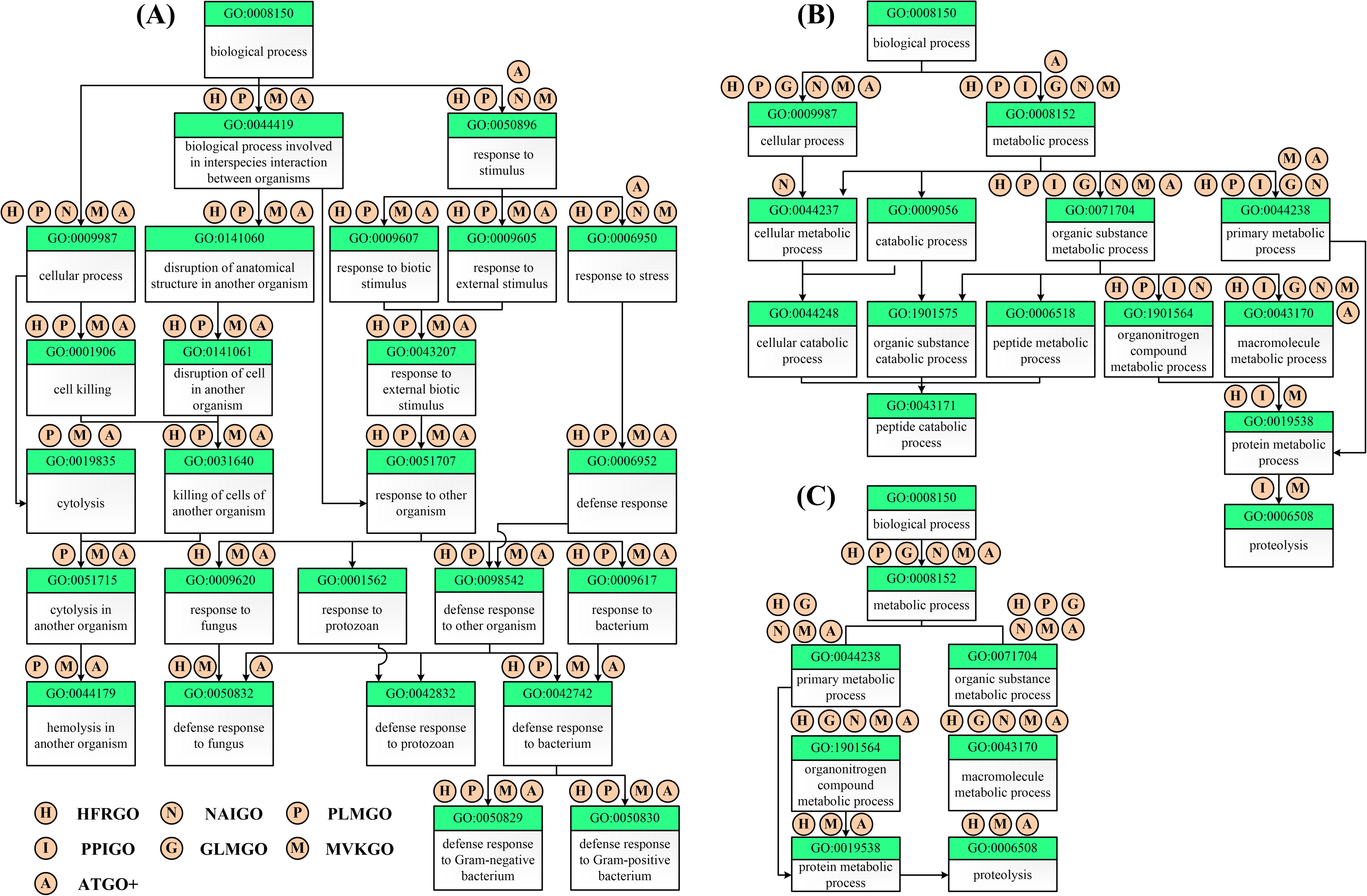
The directed acyclic graph of GO terms in BP aspect for three representative cases. The circles above each GO term represent prediction methods, where a circle filled with “X” on GO term “Y” signifies that method “X” correctly predicts term “Y”. **(A)** A0A2L2DDE6. **(B)** Q8I2J3. **(C)** Q7Q2T8.

**Table 2.** The modeling results of MKFGO in comparison with 6 competing GO prediction methods on three representative cases in BP prediction.

| Method | A0A2L2DDE6 |  |  | Q8I2J3 |  |  | Q7Q2T8 |  |  |
| --- | --- | --- | --- | --- | --- | --- | --- | --- | --- |
|  | TP | FP | F<sub>1</sub>-score | TP | FP | F<sub>1</sub>-score | TP | FP | F<sub>1</sub>-score |
| HFRGO | 20 | 5 | 0.800 | 7 | 11 | 0.438 | 7 | 5 | 0.737 |
| PLMGO | 21 | 0 | 0.913 | 5 | 1 | 0.500 | 2 | 2 | 0.364 |
| PPIGO | 0 | 0 | 0.000 | 7 | 7 | 0.500 | 0 | 0 | 0.000 |
| NAIGO | 3 | 33 | 0.098 | 7 | 29 | 0.280 | 5 | 31 | 0.233 |
| DLMGO | 0 | 0 | 0.000 | 5 | 9 | 0.357 | 5 | 18 | 0.333 |
| ATGO+ | 23 | 9 | 0.807 | 6 | 0 | 0.600 | 7 | 10 | 0.583 |
| MKFGO | 23 | 1 | **0.938** | 8 | 0 | **0.727** | 7 | 2 | **0.875** |

TP: the number of correctly predicted GO terms.
FP: the number of mistakenly predicted GO terms.
Bold fonts highlight the best performer in each category.

These data reveal several interesting insights. Overall, MKFGO is the best performer with the highest F_1_-score among all seven GO prediction methods across three cases. In A0A2L2DDE6, HFRGO and PLMGO predict nearly the same number of true positives, with 20 and 21 GO terms, respectively, significantly outperforming the other three component methods (PPIGO, NAIGO, and DLMGO). Importantly, among the five component methods, either PLMGO or HFRGO can correctly predict these 5 GO terms: GO:0051715, GO:0019835, GO:0001906, GO:0009620, and GO:0050832. After incorporating five components, MKFGO successfully inherits all of the 23 true positives. In Q8I2J3, five components gain a total of 9 true positives, where only NAIGO and PPIGO separately correctly identify GO:0044237 and GO:000605 with confidence scores of 0.208 and 1.000, respectively. As a result, the composite MKFGO yields 8 true positives without false positives. It cannot escape our notice that the GO:0044237 is excluded in the modeling results of MKFGO. The underlying reason is that the low confidence score of 0.208 from NAIGO is further diluted to 0.074, falling below the cut-off value of MKFGO, after decision-level fusion. Occasionally, the main contributors (HFRGO, PLMGO, and PPIGO) cannot provide any predicted GO terms, as observed in the case of Q9SV19, listed in **Table S7**. Even in this case, MKFGO can inherit part of predictions from NAIGO and DLMGO, maintaining acceptable performance. These cases demonstrate that the complementary functional knowledge embedded in different component methods can be effectively integrated into MKFGO.

Sometimes, one component method can capture all true positives yielded by other methods. Taking Q7Q2T8 as an example, HFRGO correctly hit all 7 GO terms, covering the true positives of the other four components. Other examples include Q9KG76, A0A1D5RMD1, and J9VWW9, in which the PLMGO, PPIGO, and DLMGO could individually encompass all true positives predicted by other components, as listed in **Tables S8, S9,** and **S10**. Even in these cases, the final MKFGO can effectively integrate all true positives from the five components with the least false positives, leading to further improved performance.

## CONCLUSION

We developed a novel composite deep learning method, MKFGO, to predict protein functions using the integration of five GO prediction pipelines built on multi-source biological data. Large-scale benchmarking on 1522 non-redundant test proteins demonstrated that MKFGO consistently outperforms 11 existing state-of-the-art methods in GO prediction accuracy. The performance advantage of MKFGO mainly stems from several advancements. First, two deep-learning component methods, HFRGO and PLMGO, could capture the functional-related knowledge from protein sequences in different views, with effective knowledge fusion at the decision level. Specifically, HFRGO derived three handcrafted features from the views of sequence conversion, secondary structure, and family domain, which are then associated with function prediction through integrating the designed LSTM-attention network with the TL-GBA strategy. PLMGO employs the ProtTrans transformer to encode the sequences into feature embeddings with evolution diversity, then decoded by the fully connected neural network. Second, another three components, driven by protein-protein interaction, GO term probability, and coding-gene sequence, provide complementary knowledge for function prediction.

Despite the promising prediction performance, there remains significant potential for further improvements. First, the confidence scores from the five component methods are merged into a consensus score using a simple one-layer fully connected neural network. However, employing a more advanced deep learning approach could further enhance the integration of confidence scores. Second, with the advancement of protein structure prediction models (e.g., AlphaFold3 [62] and ESMFold [32]), the predicted three-dimensional structures hold great potential for enhancing function prediction. Research in these areas is currently ongoing.

## ACKNOWLEDGMENTS

This work is supported by the National Natural Science Foundation of China (No. 62402227, No. 62306142 and No. 62372234), Fundamental Research Funds for the Central Universities (YDZX2025024), Jiangsu Funding Program for Excellent Postdoctoral Talent (No. 2023ZB224), Natural Science Foundation of the Higher Education Institutions of Jiangsu Province of China under grant 24KJB520041. We thank Prof. Zhiwei Ji, Dr. Fan Fei, and Dr. Xiao Han for the insightful discussions during the lab meeting.

## Key Points

- Accurate determination of protein functions is crucial for understanding life mechanisms and advancing drug discovery. This study has developed MKFGO, a novel composite deep learning model, to predict GO terms of proteins by integrating five complementary pipelines built on multi-source biological data.
- Experimental results demonstrate that MKFGO significantly outperforms existing state-of-the-art methods in GO prediction accuracy. The key strength of MKFGO lies in its two deep learning components, HFRGO and PLMGO, which extract functional knowledge from protein sequences in different views, with effective knowledge fusion at the decision level.
- HFRGO leverages an LSTM-attention network embedded with handcraft features, in which a TL-GBA strategy is designed to strengthen the correlation between feature similarity and function similarity. PLMGO utilizes the ProtTrans transformer to encode the sequences into feature embeddings with evolution diversity, which are then decoded using the fully connected neural network.

## Code availability

The source codes and models can be freely downloaded at https://github.com/yiheng-zhu/MKFGO.

## Author Biography

**Yi-Heng Zhu** received his Ph.D. degree in control science and engineering from Nanjing University of Science and Technology in 2023. He is currently a lecturer at the College of Artificial Intelligence, Nanjing Agricultural University. His research interests include bioinformatics, machine learning, and pattern recognition.

**Shuxin Zhu** is associate professor at the College of Artificial Intelligence, Nanjing Agricultural University. Her research interests include bioinformatics, machine learning, and pattern recognition. She is a member of the China Computer Federation (CCF).

**Xuan Yu** is currently a PHD student at Department of Computer Science, City University of Hong Kong. His research interests including bioinformatics, machine learning and artificial intelligence.

**He Yan** received the M.S. degree from Nanjing Forestry University, Jiangsu, China, in 2014, and the Ph.D. degree at Nanjing University of Science and Technology, China, in 2022. From 2019 to 2021, he acted as a visiting student at the University of Alberta in Canada. He is currently an Associate Professor with the computer science department at the Nanjing Forestry University, Nanjing, China. His research interests include bioinformatics, machine learning, artificial intelligence, data mining and intelligent medical systems.

**Yan Liu** received his Ph.D. degree in computer science from Nanjing University of Science and Technology in 2023. He is currently a young hundred distinguished professor at the School of Information Engineering, Yangzhou University, China. His research interests include pattern recognition, machine learning, and bioinformatics.

**Xiaojun Xie** received his Ph.D. degree in software engineering from Nanjing University of Aeronautics and Astronautics in 2020. He is currently a lecturer at the College of Artificial Intelligence, Nanjing Agricultural University. His research interests include bioinformatics, rough sets, granular computing, and machine learning.

**Dong-Jun Yu** is a full professor at the School of Computer Science and Engineering, Nanjing University of Science and Technology, Nanjing, China. His research interests include pattern recognition, machine learning, and bioinformatics. He is a senior member of the China Computer Federation (CCF) and a senior member of the China Association of Artificial Intelligence (CAAI).

**Rui Ye** received her Ph.D. degree in College of Computer Science and Technology from Nanjing University of Aeronautics and Astronautics in 2023. She is currently a lecturer in College of Artificial Intelligence, Nanjing Agricultural University. Her research interests include bioinformatics, pattern recognition, data mining and time series forecasting.

