## Supplementary Materials for "MKFGO: Integrating Multi-Source Knowledge Fusion with Pre-Trained Language Model for High-Accuracy Protein Function Prediction"

**Table of content**

**Supporting Text**

- Text S1. Feature representation.
- Text S2. The procedures for the LSTM-attention network.
- Text S3. The functional similarity between two proteins.
- Text S4. The details of the PPIGO pipeline.
- Text S5. Hierarchy of GO annotations.
- Text S6. Evaluation metrics.
- Text S7. Decision-level fusion shows superior performance than feature-level fusion.
- Text S8. Construction of gene benchmark dataset.
- Text S9. Ensemble techniques.

**Supporting Figures**

- Figure S1. The AUROC values of 12 protein function prediction methods across three GO aspects on 300 test proteins from 158 new species.
- Figure S2. The AUROC values of 11 protein function prediction methods across three GO aspects on 305 non-homology test proteins.
- Figure S3. The flowchart of CM1 incorporating the PLM-based features of PLMGO into HFRGO’s architecture through feature concatenation.
- Figure S4. The flowchart of three ablation models for HFRGO.

**Supporting Tables**

- Table S1. The numbers of proteins and GO terms in the benchmark dataset.
- Table S2. The values of $d_{m}$, $c_{f}$, $\alpha$, and $K$ for three GO aspects.
- Table S3. The performance comparison between 15 function prediction methods on a subset of 515 test proteins.
- Table S4. The performance comparison between MKFGO with three control methods on 1522 test proteins.
- Table S5. The performance comparison between three gene function prediction methods on 147 non-coding genes.
- Table S6. The false positives predicted by 8 GO prediction methods on the BP aspect for three cases.
- Table S7. The predicted GO terms by 8 GO prediction methods on the BP aspect for Q9SV19.
- Table S8. The predicted GO terms by 8 GO prediction methods on the BP aspect for Q9KG76.
- Table S9. The predicted GO terms by 8 GO prediction methods on the BP aspect for A0A1D5RMD1.
- Table S10. The predicted GO terms by 8 GO prediction methods on the BP aspect for J9VWW9.

**Text S1. Feature representation**

For the input sequence with the length $L$, the PSI-BLAST program [1] is employed to search against the Swiss-Prot database [2] with the e-value of 0.01 and iteration number of 3 for extracting the position-specific scoring matrix (PSSM), with the scale of $L\times20$, where 20 is the number of essential amino acids (A, R, D, N, C, E, Q, G, H, I, L, K, M, F, P, S, T, W, Y, V). Then, a standard logistic function was utilized to normalize the original PSSM:

$f\left( x \right)=1/(1+e^{-x})$	(S1)

where $x$ is an element in the original PSSM.

The SPOT-1D-LM software [3] is used to predict the secondary structure type for each residue in the protein sequence, which belongs to one of eight types, including alpha-helix (H), 310-helix (G), pi-helix (I), beta-strand (E),beta-bridge (B), hydrogen-bonded turn (T), bend (S) and Coil (C). Therefore, the input sequence could be encoded as a one-hot matrix with the scale of $L\times8$, according to the types of secondary structures of residues. This matrix is named as the secondary structure coding matrix (SSCM) in this work.

The InterProScan program [4] is utilized to scan against the InterPro protein signature database [5] to produce the family and domain-based binary vector (FDBV) with 45899 dimensions, which is the total number of protein families, domains, and motifs. If a protein is associated with one family/domain/motif, the corresponding value in FDBV is equal to 1.

**Text S2. The procedures for the LSTM-attention network**

**(A) Feature concatenation**

For the query sequence with the length $L$, we extract the corresponding three feature representations, including PSSM, SSCM, and FDBV, with the scales of $L\times20$, $L\times8$, and 45899-D, respectively. The PSSM and SSCM are concatenated in the residue level to generate a hybrid feature matrix $\boldsymbol{X}$:

$\boldsymbol{X}=\left[ \mathbf{PSSM, SSCM}\boldsymbol{\times}\boldsymbol{W}_{\boldsymbol{s}} \right]\in R^{L\times148}$, $\boldsymbol{W}_{\boldsymbol{s}}\in R^{8\times128}$	(S2)

where $\boldsymbol{W}_{\boldsymbol{s}}$ is a random embedding matrix.

The hybrid matrix $\boldsymbol{X}$ is fed to an LSTM-attention module to output a 1024-dimensional feature vector $\boldsymbol{v}_{1}$, consisting of a BiLSTM layer, a self-attention layer, an average-pooling layer, and a fully connected layer (${FCL}_{1}$).

**(B) BiLSTM layer**

The BiLSTM is composed of a forward LSTM and a backward LSTM, both sharing the same architecture with 128 cells but operating in reverse propagation directions. Each LSTM cell mainly consists of two states (cell state $c$ and hidden state $h$) and three gates (forget gate $f$, input gate $i$, and output gate $o$). The cell state stores signals, while the hidden state outputs signals at the current time step. The forget, input, and output gates, respectively, control the extent to which past signals are retained, current signals are incorporated, and updated signals are output. Specifically, at time-step $t$ ($t\leq L$), the above-mentioned states and gates are computed as follows:

$\boldsymbol{h}_{t}=\boldsymbol{o}_{t}\cdot tanh(\boldsymbol{c}_{t})$	(S3)

$\boldsymbol{c}_{t}=\boldsymbol{f}_{t}\cdot\boldsymbol{c}_{t-1}+\boldsymbol{i}_{t}\cdot\boldsymbol{c}_{t}^{'}$	(S4)

$\boldsymbol{c}_{t}^{'}=tanh(\boldsymbol{w}_{c}\cdot\left[ \boldsymbol{h}_{t-1},\boldsymbol{x}_{t} \right]+\boldsymbol{b}_{c})$	(S5)

$\boldsymbol{o}_{t}=\sigma(\boldsymbol{w}_{o}\cdot\left[ \boldsymbol{h}_{t-1},\boldsymbol{x}_{t} \right]+\boldsymbol{b}_{o})$	(S6)

$\boldsymbol{f}_{t}=\sigma(\boldsymbol{w}_{f}\cdot\left[ \boldsymbol{h}_{t-1},\boldsymbol{x}_{t} \right]+\boldsymbol{b}_{f})$	(S7)

$\boldsymbol{i}_{t}=\sigma(\boldsymbol{w}_{i}\cdot\left[ \boldsymbol{h}_{t-1},\boldsymbol{x}_{t} \right]+\boldsymbol{b}_{i})$	(S8)

where $\boldsymbol{x}_{t}$ is the vector in the $t$-th row of the feature matrix $\boldsymbol{X}$, acting as the input at time-step $t$ and representing the feature vector of the $t$-th residue in the protein sequence. $\boldsymbol{c}_{t-1}$ and $\boldsymbol{h}_{t-1}$ are cell and hidden states, respectively, at the time-step $t$-1, $\boldsymbol{w}_{*}$ is the weight, $\boldsymbol{b}_{*}$ is the bias, $\left[ , \right]$ is concatenation operation between two vectors, and $\sigma(\cdot)$ is the Sigmoid function.

The output of the BiLSTM layer is represented as $\boldsymbol{H}$ with the scale of $L\times256$, obtained by concatenating the hidden states of all LSTM cells across all time steps:

$\boldsymbol{H=}\left[ \begin{aligned} \boldsymbol{h}_{1} \\ \boldsymbol{h}_{2} \\ \boldsymbol{\ldots} \\ \boldsymbol{h}_{L} \end{aligned} \right]$	(S9)

**(C) Self-attention layer**

The self-attention layer comprises 8 attention heads, each executing scaled dot-product attention as follows:

$\boldsymbol{A}_{i}=SoftMax\left( \boldsymbol{M}_{i}^{Q}\cdot\frac{{{(\boldsymbol{M}}_{i}^{K})}^{T}}{\sqrt{d_{i}}} \right)\cdot\boldsymbol{M}_{i}^{V}$, $\boldsymbol{A}_{i}\in R^{L\times32}$	(S10)

$\boldsymbol{M}_{i}^{Q}=\boldsymbol{H}{\cdot\boldsymbol{W}}_{i}^{Q}$, $\boldsymbol{M}_{i}^{K}=\boldsymbol{H}{\cdot\boldsymbol{W}}_{i}^{K}$, $\boldsymbol{M}_{i}^{V}=\boldsymbol{H}{\cdot\boldsymbol{W}}_{i}^{V}$, $\boldsymbol{M}_{i}^{Q}, \boldsymbol{M}_{i}^{K}, \boldsymbol{M}_{i}^{V}\in R^{L\times32}$	(S11)

where $\boldsymbol{A}_{i}$ is an attention matrix in the $i$-th attention head, $\boldsymbol{M}_{i}^{Q}$, $\boldsymbol{M}_{i}^{K}$, and $\boldsymbol{M}_{i}^{V}$ are Query, Key, and Value matrices, respectively, $\boldsymbol{M}_{i}^{Q}{{\cdot(\boldsymbol{M}}_{i}^{K})}^{T}$ is a $L\times L$weight matrix that quantifies the positional correlation between amino acid pairs in the Query, while $d_{i}$ is a scale factor.

**(D) Average pooling and fully connected layer**

The attention matrices from all 8 heads are concatenated and passed through an average pooling layer, followed by a fully connected layer (${FCL}_{1}$) with 1024 neurons:

$\boldsymbol{A}=\left[ \boldsymbol{A}_{1},\boldsymbol{A}_{2}, \ldots{, \boldsymbol{A}}_{8} \right] \in R^{L\times256}$	(S12)

$\boldsymbol{a}=Avg (\mathbf{A})\in R^{256}$	(S13)

$\boldsymbol{v}_{1}=Relu(\boldsymbol{W}_{a}\cdot\boldsymbol{a}+\boldsymbol{b}_{a}) \in R^{1024}$	(S14)

where $Avg(\cdot)$ is the average function, and $Relu(\cdot)$ is the linear rectification function.

**(E) Output layer**

The feature vector $\boldsymbol{v}_{1}$ is concatenated with the FDBV to generate the feature embedding vector $\boldsymbol{v}_{2}$:

$\boldsymbol{v}_{2}=[\boldsymbol{v}_{1},FDBV\times\boldsymbol{W}_{\boldsymbol{f}}]\in R^{2048}$, $\boldsymbol{W}_{\boldsymbol{f}}\in R^{45899\times1024}$	(S15)

where $\boldsymbol{W}_{\boldsymbol{f}}$ is a random embedding matrix.

The embedding vector $\boldsymbol{v}_{2}$ is fed to a fully connected layer (${FCL}_{2}$) with 1024 neurons to generate another embedding vector $\boldsymbol{v}_{3}$, which is then processed by an output layer with $n$ neurons to output the confidence score vector $\boldsymbol{s}_{sig}$:

$\boldsymbol{v}_{3}=Relu(\boldsymbol{W}_{2}\cdot\boldsymbol{v}_{2}+\boldsymbol{b}_{2})\in R^{1024}$	(S16)

$\boldsymbol{s}_{sig}=\sigma(\boldsymbol{W}_{3}\cdot\boldsymbol{v}_{3}+\boldsymbol{b}_{3})\in R^{n}$	(S17)

where $n$ is the number of candidate GO terms. The values of $n$ are set to be 6858, 19687, and 2830 for MF, BP, and CC predictions.

At the same time, the triplet loss-based guilt-by-association (TN-GBA) strategy [6] is performed on the embedding vector $\boldsymbol{v}_{3}$ to produce another confidence score vector $\boldsymbol{s}_{gba}$. Finally, two confidence score vectors are weightedly combined to generate the final confidence score vector $\boldsymbol{s}_{hfr}$ for the HFRGO pipeline:

$\boldsymbol{s}_{hfr}=w\cdot\boldsymbol{s}_{sig}+(1-w)\cdot\boldsymbol{s}_{gba}$	(S18)

**Text S3. The functional similarity between two proteins**

The functional similarity of two proteins is measured by the F1-score between their GO terms:

$F1-score=2(pre\times rec)/(pre+rec)$, $pre=ns/n_{a}$, $rec=ns/n_{b}$	(S19)

where $ns$ is the number of same GO terms between two proteins, $n_{a}$ and $n_{b}$ are the numbers of GO terms for proteins $a$ and $b$, respectively.

**Text S4. The details of PPIGO pipeline**

The Blastp [1] program is utilized to hit a PPI entry $P_{e}$, which has the highest sequence identity to the query protein, against the STRING database [7]. For each PPI partner of $P_{e}$, the Blastp is employed again with the e-value of 0.1 to search the corresponding homologs from the training sequence dataset. These homology proteins are used as templates to calculate the confidence scores of GO terms for the query:

$s_{ppi}\left( {GO}_{j} \right)=\frac{\sum_{i=1}^{N} \sum_{k}^{n_{i}} ({c_{i}\cdot b}_{i,k}\cdot I_{i,k}\left( {GO}_{j} \right))}{\sum_{i=1}^{N} \sum_{k}^{n_{i}} {c_{i}\cdot b}_{i,k}}$	(S20)

where $N$ is the number of PPI partners for $P_{e}$, and $c_{i}$ is the score assigned in the STRING database as confidence of interaction between $P_{e}$ and its $i$-th partner; $n_{i}$ is the number of homologs for the $i$-th partner of $P_{e}$ using Blastp search, and $b_{i,k}$ is the bit-score between the $i$-th partner and its $k$-th homolog by Blastp.

**Text S5. Hierarchy of GO annotations**

GO annotation follows a hierarchical structure [8]. In both ground truth and predictions, if a protein is assigned a GO term, it must also be annotated with the direct parent and all ancestral terms of this term. To ensure this hierarchical relationship, we adhere to CAFA’s rule and employ a post-processing procedure [9] to refine the confidence scores for all predicted GO terms:

${s\left( {GO}_{j} \right)}_{post}$ = max ($s\left( {GO}_{j} \right), {s\left( {GO}_{j}^{1} \right)}_{post},{s\left( {GO}_{j}^{2} \right)}_{post}, \ldots, {s\left( {GO}_{j}^{M} \right)}_{post}$)	(S21)

where $s\left( {GO}_{j} \right)$ and ${s\left( {GO}_{j} \right)}_{post}$ are the confidence scores of ${GO}_{j}$ before and after post-processing, ${s\left( {GO}_{j}^{1} \right)}_{post},{s\left( {GO}_{j}^{2} \right)}_{post}, \ldots, {s\left( {GO}_{j}^{M} \right)}_{post}$ are the confidence scores of all direct child terms of ${GO}_{j}$ after post-processing. This post-processing procedure ensures that the confidence score of a GO term is larger than or equal to those of all its children.

**Text S6. Evaluation metrics**

$F_{max}$ is the highest F-score achieved across all confidence thresholds, offering a single measure of the best trade-off between precision and recall, with the following definition:

$F_{max}=\max_{t} \{\frac{2\cdot Pre\left( t \right)\cdot Rec\left( t \right)}{Pre\left( t \right)+Rec\left( t \right)}\}$	(S22)

$Pre\left( t \right)=\frac{1}{m(t)}\cdot\sum_{i=1}^{m(t)} \frac{\sum_{j=1}^{n} 1(s_{ij}\geq t)\cdot I_{ij}}{\sum_{j=1}^{n} 1(s_{ij}\geq t)}$	(S23)

$Rec\left( t \right)=\frac{1}{m}\cdot\sum_{i=1}^{n} \frac{\sum_{j=1}^{n} 1(s_{ij}\geq t)\cdot I_{ij}}{\sum_{j=1}^{n} I_{ij}}$	(S24)

where $t$ is a threshold ranging from 0 to 1, $Pre\left( t \right)$ and $Rec\left( t \right)$ are precision and recall under the threshold $t$, respectively; $s_{ij}$ is the confidence score that the *i*-th protein is associated with the *j*-th GO term using the function prediction model; $1\left( \cdot\right)=1$, if the input is true; otherwise, $1\left( \cdot\right)=0$; $I_{ij}=1$, if the *i*-th protein is associated with the *j*-th GO term in the experimental annotations; otherwise, $I_{ij}=0$; $m(t)$ is the number of proteins which have at least one GO term with the confidence score higher than $t$; $m$ and $n$ are the number of all test proteins and GO terms, respectively.

$S_{min}$ measures the discrepancy between predicted and true GO terms by calculating the semantic distance in the GO hierarchy structure, defined as follows:

$S_{min}=\min_{t} \{\sqrt{{ru(t)}^{2}+{mi(t)}^{2}}\}$	(S25)

$ru\left( t \right)=\frac{1}{m}\cdot\sum_{i=1}^{m} \sum_{j=1}^{n} ic\left( j \right)\cdot1(s_{ij}<t)\cdot I_{ij}$	(S26)

$mi\left( t \right)=\frac{1}{m}\cdot\sum_{i=1}^{m} \sum_{j=1}^{n} ic\left( j \right)\cdot1\left( s_{ij}\geq t \right)\cdot(1-I_{ij})$	(S27)

$ic\left( j \right)={log}_{2}\frac{1}{p(j|parent(j))}$	(S28)

where $ru(t)$ and $mi(t)$ are remaining uncertainty and misinformation under the threshold $t$, respectively; $ic\left( j \right)$ is information content of the *j*-th GO term, and $p(j|parent(j))$ is the conditional probability of the *j*-th term given its parent terms within the hierarchical GO structure. Further details are provided in reference [10].

**Text S7. Decision-level fusion shows superior performance than feature-level fusion**

We designed three control methods, represented as CM1, CM2, and CM3, as follows:

- **CM1**: The combination of HFRGO and PLMGO at the feature-level. Specifically, we directly incorporated the PLM-based features of PLMGO into HFRGO’s architecture through feature concatenation, as shown in **Figure S3**.
- **CM2**: The combination of HFRGO and PLMGO at the decision-level. For each GO term, the corresponding confidence scores predicted by HFRGO and PLMGO are ensembled as a consensus score using the neural network, consisting of a full network layer with 512 neurons and an output layer with 1 neuron.
- **CM3**: The combination of CM1 and the other three components of MKFGO at the decision-level. The prediction results of M1, PPIGO, NAIGO, and GLMGO are ensembled using the same fully connected neural network in the CM2.

We further benchmarked MKFGO with the above-three control methods on all 1522 test proteins, as illustrated in **Table S4**. It could be observed that CM2 consistently outperforms CM1 for three metrics on all GO aspects. Meanwhile, the performance of CM1 in terms of F_max_, S_min_, and AUPRC on CC aspect in **Table S4** is even inferior to that of PLMGO in **Table 1**. Moreover, after integrating with the other three components (i.e., PPIGO, NAIGO, and GLMGO), the CM3 shows inferior performance to MKFGO across all evaluation metrics, except for the S_min_ value on BP aspect. These data demonstrate that the knowledge embedded in handcrafted and PLM-based features can be more effectively fused at the decision level rather than the feature-level.

**Text S8. Construction of gene benchmark dataset**

We downloaded all 84539 genes with GO annotations from National Center for Biotechnology Information (NCBI) [11], using eight experimental evidence codes, including EXP, IDA, IPI, IMP, IGI, IEP, TAS, and IC [12, 13]. We selected all of the genes from five species (i.e., Human, Mouse, Fly, Arabidopsis, and Budding Yeast) that have the available expression data in COXPRESdb [14] or ATTED-II [15] database. After this, we collected 46375 genes, including 13556 (13497 protein-coding and 59 non-coding) genes from Human species, 10799 (10720 protein-coding and 79 non-coding) genes from Mouse, 5870 (5844 protein-coding and 26 non-coding) genes from Fly, 11065 (11023 protein-coding and 42 non-coding) genes from Arabidopsis, and 5085 (5000 protein-coding and 85 non-coding) genes from Budding Yeast. For each species, the protein-coding genes are used as the training set, and the non-coding genes are equally split into the validation and test datasets. As a result, we gathered a training dataset of 46084 protein-coding genes, a validation dataset of 144 non-coding genes, and a test dataset of 147 non-coding genes.

**Text S9. Ensemble techniques**

For a GO term (${GO}_{j}$), the fully connected neural network (FCNN) incorporates the corresponding confidence scores predicted by all component methods into a consensus score, as follows:

$\boldsymbol{v}_{1}^{*}=Relu(\boldsymbol{W}_{1}^{*}\cdot\boldsymbol{s}_{j}+\boldsymbol{b}_{1}^{*})$, $\boldsymbol{s}_{j}=[{S\left( {GO}_{j} \right)}_{1}, {S\left( {GO}_{j} \right)}_{2}, \ldots, {S({GO}_{j})}_{n}]\in R^{n}$	(S29)

${S({GO}_{j})}_{FCNN}=\sigma(\boldsymbol{W}_{2}^{*}\cdot\boldsymbol{v}_{1}^{*}+\boldsymbol{b}_{2}^{*})\in R^{1}$	(S30)

where $\boldsymbol{W}_{1}^{*}\in R^{m\times n}$ and $\boldsymbol{W}_{2}^{*}\in R^{1\times m}$ are weight matrices, $\boldsymbol{b}_{1}^{*}\in R^{m}$ and $\boldsymbol{b}_{2}^{*}\in R^{1}$ are biases; ${S({GO}_{j})}_{i}$ is the confidence score of ${GO}_{j}$ predicted by the $i$-th component method; $m$ is the number of neurons, $n$ is the number of component methods. Here, $m=512$, $n=5$.

Weighted voting (WV) linearly combines the confidence scores of component methods by assigning weights to each component, as follows:

${S({GO}_{j})}_{WV}=\frac{\sum_{i=1}^{n} {w_{i}\times S({GO}_{j})}_{i}}{\sum_{i=1}^{n} w_{i}}$	(S31)

where $w_{i}$ is the weight assigned by the $i$-th component, set to be its AUPRC value on the validation dataset of 974 proteins.

Logistic regression (LR) is an improved version of WV through adding sigmoid function, as follows:

${S({GO}_{j})}_{LR}=\frac{1}{1+exp(-\sum_{i=1}^{n} {w_{i}\times S({GO}_{j})}_{i}+w_{o})}$	(S32)

where $w_{*}$ is a machine-learning parameter optimized on the validation dataset.

Weighted Product (WP) combines the confidence scores through successive multiplication, as follows:

${S({GO}_{j})}_{WP}=1-\prod_{i=1}^{n} (1-{S({GO}_{j})}_{i})$	(S33)

Table S1. The numbers of proteins and GO terms in the benchmark dataset.

| Benchmark dataset |  | $N_{MF}^{P}$ | $N_{BP}^{P}$ | $N_{CC}^{P}$ | $N_{ALL}^{P}$ |  | $N_{MF}^{T}$ | $N_{BP}^{T}$ | $N_{CC}^{T}$ | $N_{ALL}^{T}$ |
| --- | --- | --- | --- | --- | --- | --- | --- | --- | --- | --- |
| Training dataset |  | 52594 | 52977 | 45556 | 70212 |  | 6858 | 19687 | 2830 | 29375 |
| Validation dataset |  | 680 | 769 | 480 | 974 |  | 734 | 2659 | 304 | 3697 |
| Test dataset |  | 961 | 1172 | 767 | 1522 |  | 1009 | 3605 | 419 | 5033 |

$N_{MF}^{P}$/$N_{BP}^{P}$/$N_{CC}^{P}$/$N_{ALL}^{P}$: The number of proteins for MF/BP/CC/all-three aspects.

$N_{MF}^{T}$/$N_{BP}^{T}$/$N_{CC}^{T}$/$N_{ALL}^{T}$: The number of GO terms for MF/BP/CC/all-three aspects.

Table S2. The values of $d_{m}$, $c_{f}$, $\alpha$, and $K$ for three GO aspects

| GO aspect | $d_{m}$ | $c_{f}$ | $\alpha$ | $K$ |
| --- | --- | --- | --- | --- |
| MF | 0.1 | 0.8 | 0.1 | 30 |
| BP | 0.1 | 0.8 | 0.1 | 100 |
| CC | 0.1 | 0.8 | 0.1 | 100 |

Table S3. The performance comparison between 15 function prediction methods

on a subset of 515 test proteins

|  | Method | F_max_ | | |  | S_min_ | | |  | AUPRC | | |
| --- | --- | --- | --- | --- | --- | --- | --- | --- | --- | --- | --- | --- |
|  |  | MF | BP | CC |  | MF | BP | CC |  | MF | BP | CC |
| Single method | Blast-KNN ^a, d^ | 0.661 | 0.411 | 0.582 |  | 8.20 | 29.17 | 8.53 |  | 0.506 | 0.297 | 0.416 |
|  | FunFams ^a, d^ | 0.605 | 0.383 | 0.546 |  | 10.11 | 30.45 | 8.90 |  | 0.468 | 0.248 | 0.384 |
|  | PPIGO ^a, d^ | 0.452 | 0.362 | 0.597 |  | 11.72 | 30.04 | 7.97 |  | 0.250 | 0.223 | 0.387 |
|  | DeepGOCNN ^b, e^ | 0.470 | 0.316 | 0.531 |  | 11.23 | 30.97 | 9.34 |  | 0.405 | 0.228 | 0.531 |
|  | TALE ^b, d^ | 0.466 | 0.335 | 0.574 |  | 11.52 | 30.26 | 8.46 |  | 0.414 | 0.246 | 0.577 |
|  | DeepGOZero ^b, d^ | 0.677 | 0.391 | 0.567 |  | 8.11 | 29.76 | 9.32 |  | 0.665 | 0.313 | 0.558 |
|  | AnnoPRO ^b, e^ | 0.566 | 0.386 | 0.583 |  | 11.24 | 29.64 | 8.13 |  | 0.445 | 0.291 | 0.594 |
|  | HFRGO ^b^ | 0.690 | 0.421 | 0.611 |  | 7.56 | 28.13 | 8.17 |  | 0.655 | 0.344 | 0.566 |
|  | ATGO ^c, d^ | 0.693 | 0.429 | 0.625 |  | 7.62 | 28.27 | 8.02 |  | 0.707 | 0.364 | 0.651 |
|  | DeepGO-SE ^c, d^ | 0.674 | 0.399 | 0.590 |  | 7.97 | 29.68 | 9.27 |  | 0.694 | 0.330 | 0.622 |
|  | PLMGO ^c^ | 0.680 | 0.424 | 0.642 |  | 8.05 | 28.57 | 7.69 |  | 0.673 | 0.349 | 0.595 |
| Composite method | DeepGOPlus ^d^ | 0.647 | 0.403 | 0.576 |  | 8.15 | 29.23 | 8.40 |  | 0.636 | 0.328 | 0.588 |
|  | TALE+ ^d^ | 0.640 | 0.413 | 0.603 |  | 8.42 | 29.17 | 8.19 |  | 0.634 | 0.334 | 0.613 |
|  | ATGO+ ^d^ | 0.692 | 0.437 | 0.622 |  | 7.56 | 28.23 | 8.18 |  | 0.700 | 0.372 | 0.645 |
|  | MKFGO | **0.711** | **0.462** | **0.656** |  | **7.36** | **27.24** | **7.42** |  | **0.727** | **0.405** | **0.702** |

^a^ Template detection-based methods; ^b^ Deep learning-based methods with handcraft feature representations; ^c^ Deep learning-based methods with PLM-based feature representations; ^d^ The prediction models are re-trained on our training dataset using the author’s source codes; ^e^ The prediction models are directly downloaded from author’s web platforms. Bold fonts highlight the best performer in each category.

Table S4. The performance comparison between MKFGO with

three control methods on 1522 test proteins

| Method | F_max_ | | |  | S_min_ | | |  | AUPRC | | |
| --- | --- | --- | --- | --- | --- | --- | --- | --- | --- | --- | --- |
|  | MF | BP | CC |  | MF | BP | CC |  | MF | BP | CC |
| CM1 | 0.690 | 0.433 | 0.605 |  | 7.20 | 23.55 | 7.87 |  | 0.660 | 0.371 | 0.558 |
| CM2 | 0.700 | 0.438 | 0.630 |  | 7.02 | 23.39 | 7.57 |  | 0.706 | 0.379 | 0.598 |
| CM3 | 0.692 | 0.451 | 0.613 |  | 7.19 | **23.07** | 7.74 |  | 0.690 | 0.388 | 0.647 |
| MKFGO | **0.710** | **0.459** | **0.639** |  | **6.97** | 23.08 | **7.38** |  | **0.716** | **0.400** | **0.668** |

Bold fonts highlight the best performer in each category.

Table S5. The performance comparison between three gene function prediction methods

on 147 non-coding genes

| Method | F_max_ | | |  | S_min_ | | |  | AUPRC | | |
| --- | --- | --- | --- | --- | --- | --- | --- | --- | --- | --- | --- |
|  | MF | BP | CC |  | MF | BP | CC |  | MF | BP | CC |
| GLMGO | 0.514 | 0.344 | 0.421 |  | 8.58 | 18.08 | 11.53 |  | 0.399 | 0.258 | 0.316 |
| TripletGO ^a^ | 0.569 | 0.383 | 0.434 |  | 8.48 | 18.03 | 11.40 |  | 0.424 | 0.289 | 0.310 |
| TripletGO + GLMGO ^b^ | **0.588** | **0.399** | **0.473** |  | **8.42** | **17.82** | **10.81** |  | **0.463** | **0.305** | **0.390** |

^a^ The prediction models are re-trained on our training dataset using the source code. ^b^ The composite method utilizes the fully connected neural network to ensemble the TripletGO and GLMGO prediction results. Bold fonts highlight the best performer in each category.

Table S6. The false positives predicted by 8 GO prediction methods on the BP aspect for three cases

| Case | Method | False Positive |
| --- | --- | --- |
| A0A2L2DDE6 | HFRGO | GO:0050789 GO:0140546 GO:0006955 GO:0065007 GO:0002376 |
|  | PLMGO |  |
|  | PPIGO |  |
|  | NAIGO | GO:0044237 GO:0050789 GO:0051234 GO:0019219 GO:0009889 GO:0060255 GO:0071704 GO:0031326 GO:0048523 GO:1901360 GO:0048519 GO:1901576 GO:0065007 GO:0032501 GO:0080090 GO:0008152 GO:0031323 GO:0016043 GO:0010468 GO:0050794 GO:0071840 GO:0032502 GO:0048856 GO:0048518 GO:0006810 GO:0051179 GO:0019222 GO:0043170 GO:0048522 GO:0044238 GO:1901564 GO:0010556 GO:0009058 |
|  | GLMGO |  |
|  | ATGO+ | GO:0044092 GO:0050789 GO:0043086 GO:0050790 GO:0002376 GO:0051239 GO:0065007 GO:0032501 GO:0065009 |
|  | MKFGO | GO:0002376 |
| A0A2L2DDE6 | HFRGO | GO:0006810 GO:0071705 GO:0045184 GO:0051641 GO:0051179 GO:0071702 GO:0051234 GO:0008104 GO:0070727 GO:0015031 GO:0033036 |
|  | PLMGO | GO:1901360 |
|  | PPIGO | GO:0044129 GO:0050789 GO:0044145 GO:0044127 GO:0044149 GO:0065007 GO:0048518 |
|  | NAIGO | GO:0050789 GO:0051234 GO:0009889 GO:0019219 GO:0060255 GO:0050896 GO:0031326 GO:0048523 GO:1901360 GO:0048519 GO:0006950 GO:1901576 GO:0065007 GO:0032501 GO:0080090 GO:0031323 GO:0016043 GO:0010468 GO:0050794 GO:0071840 GO:0032502 GO:0048856 GO:0048518 GO:0006810 GO:0051179 GO:0019222 GO:0048522 GO:0010556 GO:0009058 |
|  | GLMGO | GO:1901360 GO:0016043 GO:0050789 GO:0090304 GO:0006139 GO:0050794 GO:0050896 GO:0071840 GO:0065007 |
|  | ATGO+ |  |
|  | MKFGO |  |
| Q7Q2T8 | HFRGO | GO:0009987 GO:0050789 GO:1901575 GO:0009056 GO:0065007 |
|  | PLMGO | GO:0009987 GO:0032501 |
|  | PPIGO |  |
|  | NAIGO | GO:0044237 GO:0050789 GO:0051234 GO:0009889 GO:0019219 GO:0060255 GO:0050896 GO:0031326 GO:0048523 GO:1901360 GO:0048519 GO:0006950 GO:1901576 GO:0065007 GO:0032501 GO:0080090 GO:0031323 GO:0016043 GO:0010468 GO:0050794 GO:0071840 GO:0032502 GO:0048856 GO:0048518 GO:0006810 GO:0051179 GO:0019222 GO:0048522 GO:0009987 GO:0010556 GO:0009058 |
|  | GLMGO | GO:0016052 GO:1901360 GO:0044237 GO:0044249 GO:0009987 GO:0006950 GO:1901575 GO:0005976 GO:0044419 GO:0044281 GO:0071840 GO:0009056 GO:0050896 GO:1901576 GO:0065007 GO:0005975 GO:0006629 GO:0009058 |
|  | ATGO+ | GO:0009987 GO:0050789 GO:1901575 GO:0065007 GO:0051239 GO:0009056 GO:0050794 GO:0032502 GO:0048856 GO:0032501 |
|  | MKFGO | GO:0009987 GO:0009056 |

Table S7. The predicted GO terms by 8 GO prediction methods on the BP aspect for Q9SV19

| Method | Predicted GO terms |
| --- | --- |
| Ground truth | GO:0006950 GO:0006979 GO:0010035 GO:0010038 GO:0010043 GO:0042221 GO:0046686 GO:0050896 (**8 terms**) |
| HFRGO |  |
| PLMGO |  |
| PPIGO |  |
| NAIGO | GO:0006950 GO:0050896 GO:0044237 GO:0050789 GO:0051234 GO:0009889 GO:0019219 GO:0060255 GO:0071704 GO:0031326 GO:0048523 GO:1901360 GO:0048519 GO:1901576 GO:0065007 GO:0032501 GO:0080090 GO:0008152 GO:0031323 GO:0016043 GO:0010468 GO:0050794 GO:0071840 GO:0032502 GO:0048856 GO:0048518 GO:0006810 GO:0051179 GO:0019222 GO:0043170 GO:0048522 GO:0009987 GO:0044238 GO:1901564 GO:0010556 GO:0009058  **TP=2, FP=34, F_1_=0.091** |
| GLMGO | GO:0006950 GO:0042221 GO:0050896 GO:0009755 GO:2001141 GO:0050789 GO:0009719 GO:0022414 GO:0019219 GO:0009889 GO:0007165 GO:0060255 GO:0009893 GO:0006355 GO:0031326 GO:0048523 GO:0048519 GO:0065007 GO:0009628 GO:0032501 GO:0080090 GO:0010033 GO:0031323 GO:0009725 GO:0010468 GO:0048583 GO:1901700 GO:0050794 GO:0032502 GO:0050793 GO:0048856 GO:0048518 GO:0019222 GO:0048522 GO:0009987 GO:0051252 GO:0003006 GO:0010556 GO:0033993 GO:0099402  **TP=3, FP=37, F_1_=0.125** |
| ATGO+ | GO:0050789 GO:0065007 GO:0050794  **TP=0, FP=3, F_1_=0.000** |
| MKFGO | GO:0050896 GO:0019222 GO:0031323 GO:0050789 GO:0009987 GO:0065007 GO:0060255 GO:0050794 GO:0032502 GO:0080090 |
|  | **TP=1, FP=9, F_1_=0.111** |

Table S8. The predicted GO terms by 8 GO prediction methods on the BP aspect for Q9KG76

| Method | Predicted GO terms |
| --- | --- |
| Ground truth | GO:0000272 GO:0005975 GO:0005976 GO:0008152 GO:0009056 GO:0009057 GO:0016052 GO:0043170 GO:0044238 GO:0071704 GO:1901575 (**11 terms**) |
| HFRGO | GO:0016043 GO:0009987 GO:0050789 GO:0044419 GO:0071840 GO:0065007  **TP=0, FP=6, F_1_=0.000** |
| PLMGO | GO:0000272 GO:0005975 GO:0005976 GO:0008152 GO:0009056 GO:0009057 GO:0016052 GO:0043170 GO:0044238 GO:0071704 GO:1901575 GO:0010383 GO:0045491 GO:0009987 GO:0016998 GO:1901564 GO:0010410 GO:0044036 GO:0045493 GO:2000895 GO:0044347  **TP=11, FP=10, F_1_=0.688** |
| PPIGO |  |
| NAIGO | GO:0008152 GO:0043170 GO:0044238 GO:0071704 GO:0044237 GO:0050789 GO:0051234 GO:0019219 GO:0009889 GO:0060255 GO:0050896 GO:0031326 GO:0048523 GO:1901360 GO:0048519 GO:0006950 GO:1901576 GO:0065007 GO:0032501 GO:0080090 GO:0031323 GO:0016043 GO:0010468 GO:0050794 GO:0071840 GO:0032502 GO:0048856 GO:0048518 GO:0006810 GO:0051179 GO:0019222 GO:0048522 GO:0009987 GO:1901564 GO:0010556 GO:0009058  **TP=4, FP=32, F_1_=0.170** |
| GLMGO |  |
| ATGO+ | GO:0000272 GO:0005975 GO:0005976 GO:0008152 GO:0009056 GO:0009057 GO:0016052 GO:0043170 GO:0044238 GO:0071704 GO:1901575 GO:0010383 GO:0009987 GO:0045491 GO:0016998 GO:0010410 GO:0044036 GO:0045493 GO:2000895 GO:0044347  **TP=11, FP=9, F_1_=0.710** |
| MKFGO | GO:0000272 GO:0005975 GO:0005976 GO:0008152 GO:0009056 GO:0009057 GO:0016052 GO:0043170 GO:0044238 GO:0071704 GO:1901575 GO:0009987 |
|  | **TP=11, FP=1, F_1_=0.957** |

Table S9. The predicted GO terms by 8 GO prediction methods on the BP aspect for A0A1D5RMD1

| Method | Predicted GO terms |
| --- | --- |
| Ground truth | GO:0003006 GO:0007281 GO:0007286 GO:0009987 GO:0022412 GO:0022414 GO:0032502 GO:0048468 GO:0048609 GO:0048856 GO:0048869 (**11 terms**) |
| HFRGO | GO:0003006 GO:0009987 GO:0022414 GO:0032502 GO:0048856 GO:0048869  **TP=6, FP=0, F_1_=0.701** |
| PLMGO | GO:0009987 GO:0016043 GO:0050789 GO:0071840 GO:0050794 GO:0065007  **TP=1, FP=5, F_1_=0.118** |
| PPIGO | GO:0003006 GO:0007281 GO:0007286 GO:0009987 GO:0022412 GO:0022414 GO:0032502 GO:0048468 GO:0048609 GO:0048856 GO:0048869 GO:0003341 GO:0060294 GO:0030317 GO:0097722 GO:0048870 GO:0007017 GO:0007018 GO:0060285 GO:0001539  **TP=11, FP=9, F_1_=0.710** |
| NAIGO | GO:0009987 GO:0032502 GO:0048856 GO:0044237 GO:0050789 GO:0051234 GO:0019219 GO:0009889 GO:0060255 GO:0050896 GO:0071704 GO:0031326 GO:0048523 GO:1901360 GO:0048519 GO:0006950 GO:1901576 GO:0065007 GO:0032501 GO:0080090 GO:0008152 GO:0031323 GO:0016043 GO:0010468 GO:0050794 GO:0071840 GO:0048518 GO:0006810 GO:0051179 GO:0019222 GO:0043170 GO:0048522 GO:0044238 GO:1901564 GO:0010556 GO:0009058  **TP=3, FP=33, F_1_=0.128** |
| GLMGO |  |
| ATGO+ | GO:0003006 GO:0007281 GO:0007286 GO:0009987 GO:0022412 GO:0022414 GO:0032502 GO:0048468 GO:0048609 GO:0048856 GO:0048869 GO:0051179 GO:0016043 GO:0050789 GO:0071840 GO:0050794 GO:0065007  **TP=11, FP=6, F_1_=0.789** |
| MKFGO | GO:0003006 GO:0007281 GO:0007286 GO:0009987 GO:0022412 GO:0022414 GO:0032502 GO:0048468 GO:0048609 GO:0048856 GO:0048869 GO:0007017 GO:0065007  **TP=11, FP=2, F_1_=0.917** |

Table S10. The predicted GO terms by 8 GO prediction methods on the BP aspect for J9VWW9

| Method | Predicted GO terms |
| --- | --- |
| Ground truth | GO:0006801 GO:0006950 GO:0006979 GO:0008152 GO:0009605 GO:0009607 GO:0009987 GO:0019430 GO:0033554 GO:0034599 GO:0035821 GO:0042221 GO:0043207 GO:0044003 GO:0044237 GO:0044403 GO:0044419 GO:0050896 GO:0051701 GO:0051707 GO:0051716 GO:0052031 GO:0052164 GO:0052167 GO:0052173 GO:0052200 GO:0052553 GO:0052572 GO:0062197 GO:0070887 GO:0072593 GO:0075136 GO:0098754 GO:0098869 GO:1990748  (**35 terms**) |
| HFRGO | GO:0006950 GO:0042221 GO:0050896 GO:0009737 GO:0006970 GO:0010033 GO:0009725 GO:0009719 GO:1901700 GO:0033993 GO:0097305 GO:0009628  **TP=3, FP=9, F_1_=0.128** |
| PLMGO | GO:0006950 GO:0008152 GO:0009987 GO:0044237 GO:0050896 GO:0071704 GO:0044238  **TP=5, FP=2, F_1_=0.238** |
| PPIGO |  |
| NAIGO | GO:0006950 GO:0008152 GO:0009987 GO:0044237 GO:0050896 GO:0050789 GO:0051234 GO:0009889 GO:0019219 GO:0060255 GO:0071704 GO:0031326 GO:0048523 GO:1901360 GO:0048519 GO:1901576 GO:0065007 GO:0032501 GO:0080090 GO:0031323 GO:0016043 GO:0010468 GO:0050794 GO:0071840 GO:0032502 GO:0048856 GO:0048518 GO:0006810 GO:0051179 GO:0019222 GO:0043170 GO:0048522 GO:0044238 GO:1901564 GO:0010556 GO:0009058  **TP=5, FP=31, F_1_=0.141** |
| GLMGO | GO:0006950 GO:0008152 GO:0009987 GO:0042221 GO:0044237 GO:0050896 GO:0071704 GO:0044249 GO:0019752 GO:1901564 GO:0044238 GO:0044281 GO:0006082 GO:0009056 GO:0009058 GO:1901576 GO:0032502 GO:0009628 GO:0043436  **TP=6, FP=13, F_1_=0.222** |
| ATGO+ | GO:0006950 GO:0008152 GO:0009987 GO:0042221 GO:0044237 GO:0050896 GO:0006970 GO:0010033 GO:0009725 GO:0009719 GO:1901700 GO:0033993 GO:0097305 GO:0009628  **TP=6, FP=8, F_1_=0.245** |
| MKFGO | GO:0006950 GO:0008152 GO:0009987 GO:0042221 GO:0044237 GO:0050896  GO:0071704 GO:0010033 GO:1901700 GO:0097305 GO:0009628  **TP=6, FP=5, F_1_=0.261** |

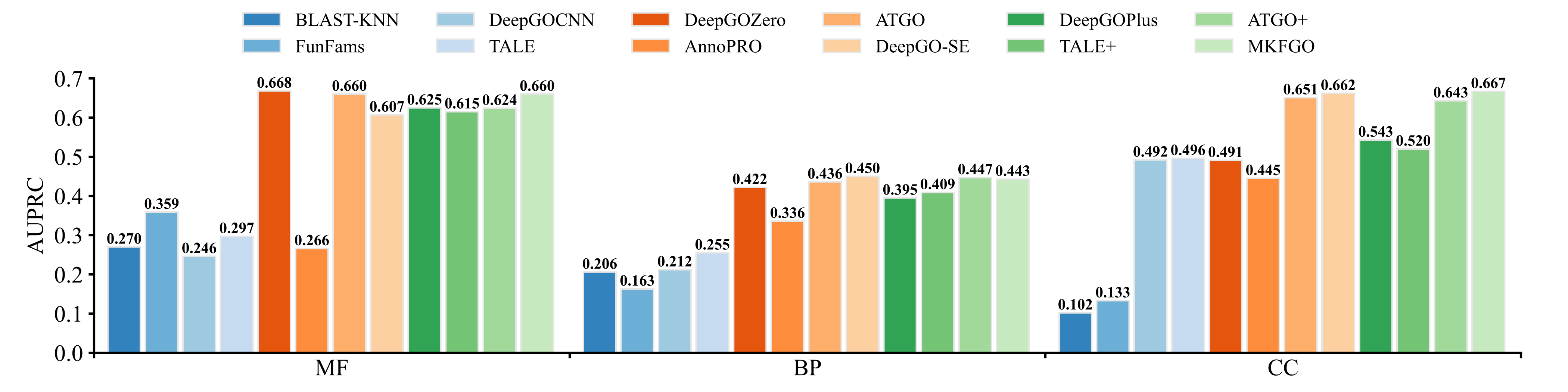
Figure S1. The AUROC values of 12 protein function prediction methods across three GO aspects on 300 test proteins from 158 new species.

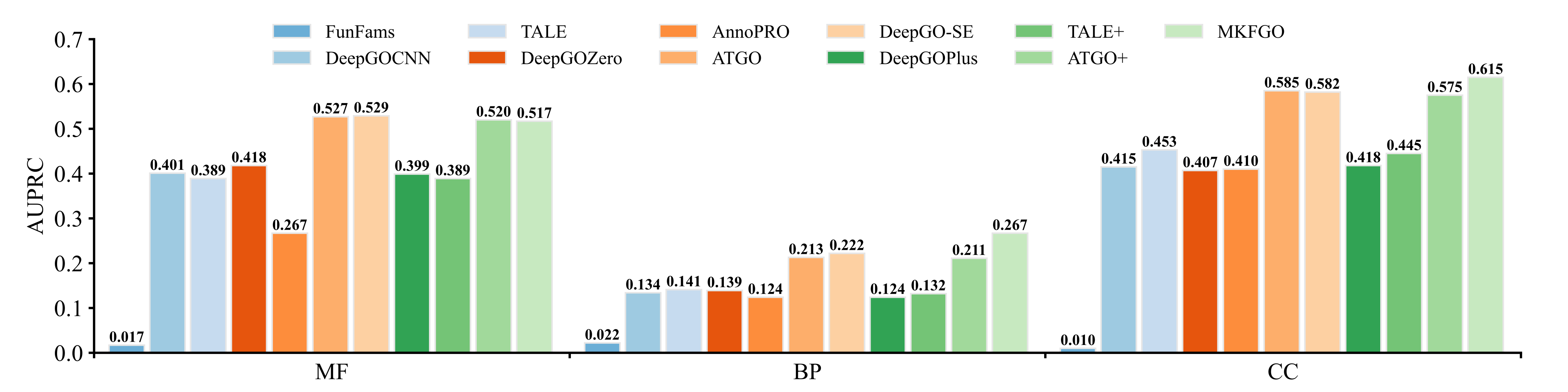

Figure S2. The AUROC values of 11 protein function prediction methods across three GO aspects on 305 non-homology test proteins.

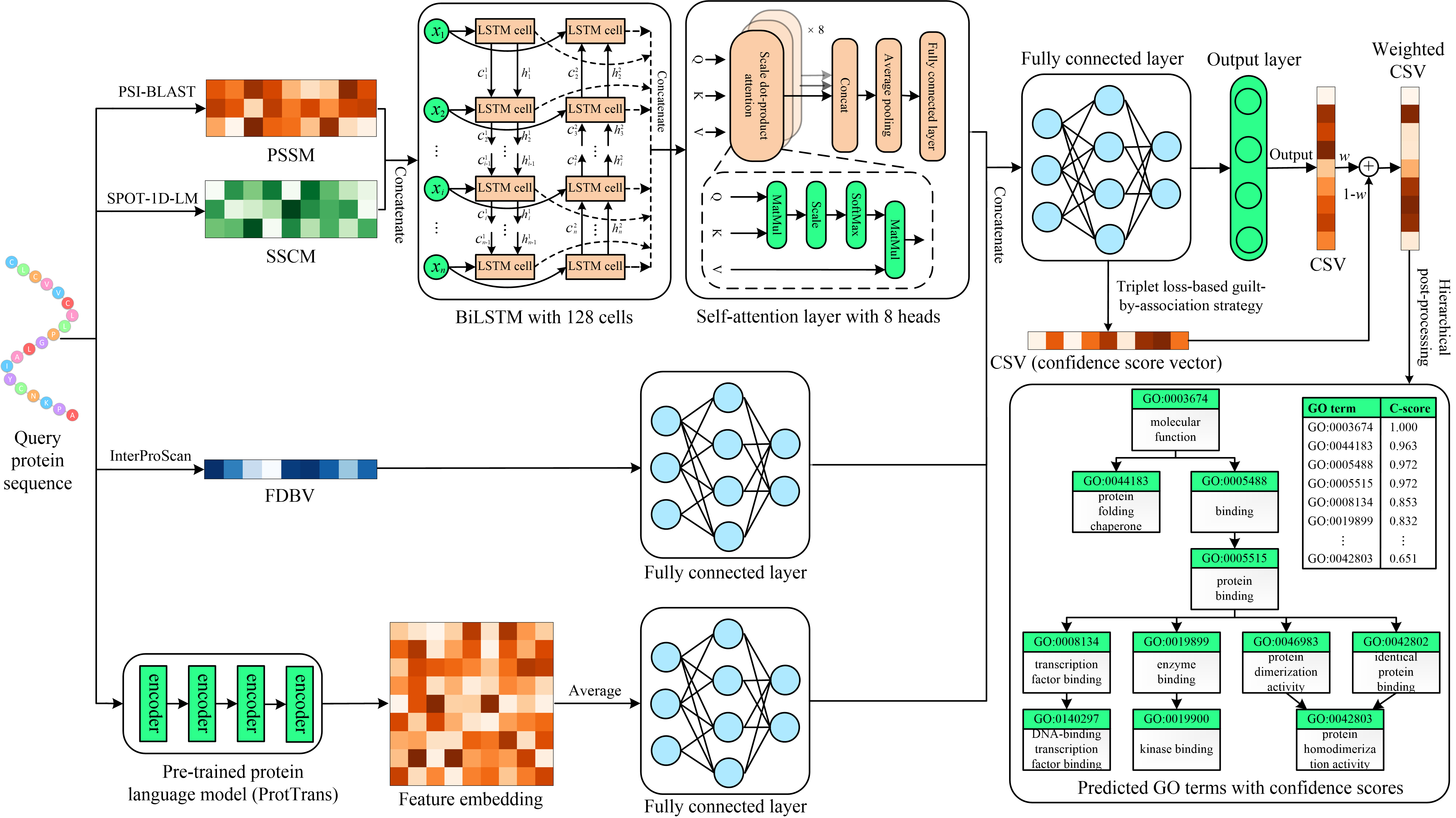

Figure S3. The flowchart of CM1 incorporating the PLM-based features of PLMGO into HFRGO’s architecture through feature concatenation.

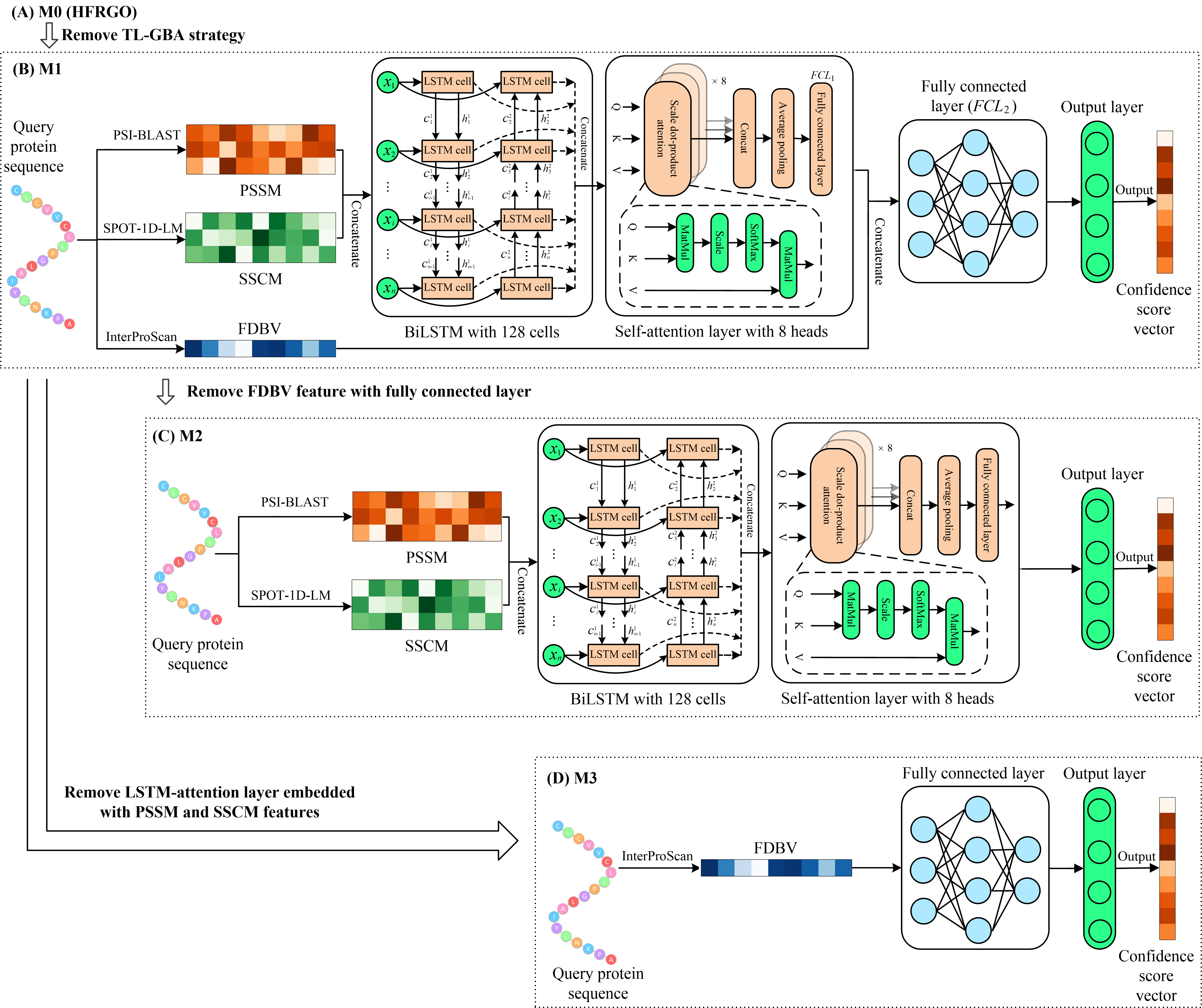

Figure S4. The flowchart of three ablation models for HFRGO.
